# Use of High Pressure NMR Spectroscopy to Rapidly Identify Proteins with Internal Ligand-Binding Voids

**DOI:** 10.1101/2020.08.25.267195

**Authors:** Donald Gagné, Roksana Azad, James M. Aramini, Xingjian Xu, Eta A. Isiorho, Uthama R. Edupuganti, Justin P. Williams, Leandro Pimentel Marcelino, Bruce A. Johnson, Kazuyuki Akasaka, Kevin H. Gardner

## Abstract

Small molecule binding within internal cavities provides a way to control protein function and structure, as exhibited in numerous natural and artificial settings. Unfortunately, most ways to identify suitable cavities require high-resolution structures *a priori* and may miss potential sites. Here we address this limitation via high-pressure solution NMR spectroscopy, taking advantage of the distinctive nonlinear pressure-induced chemical shift changes observed in proteins containing internal cavities and voids. We developed a method to rapidly characterize such nonlinearity among backbone ^1^H and ^15^N amide signals without needing to have sequence-specific chemical shift assignments, taking advantage of routinely available ^15^N-labeled samples, instrumentation, and 2D ^1^H/^15^N HSQC experiments. From such data, we find a strong correlation in the site-to-site variability in such nonlinearity with the total void volume within proteins, providing insights useful for prioritizing domains for ligand binding and indicating mode-of-action among such protein/ligand systems. We suggest that this experimental approach is a rapid and useful probe of otherwise hidden dynamic architectures of proteins, providing novel insights and opportunities into ligand binding and control.

**Significance Statement:** Many proteins can be regulated by internally binding small molecule ligands, but it is often not clear *a priori* which proteins are controllable in such a way. Here we describe a rapid method to address this challenge, using solution NMR spectroscopy to monitor the response of proteins to the application of high pressure. While the locations of NMR signals from many proteins respond to high pressure with linear chemical shift changes, proteins containing internal cavities that can bind small molecule ligands respond with easily identified non-linear changes. We demonstrate this approach on several proteins and protein/ligand complexes, suggesting that it has general utility.

## Introduction

Small molecule cofactors and ligands play critical roles in controlling protein structure and function, the understanding of which often gives insight both into natural and artificial modes of regulation. Of particular interest are identifying sites where the irregular structures of proteins give rise to cavities, voids and other features which can serve as internal sites for such compounds to bind (**Fig. 1**) and allosterically control protein structure and function (1, 6, 11–13).

**Fig. 1:**
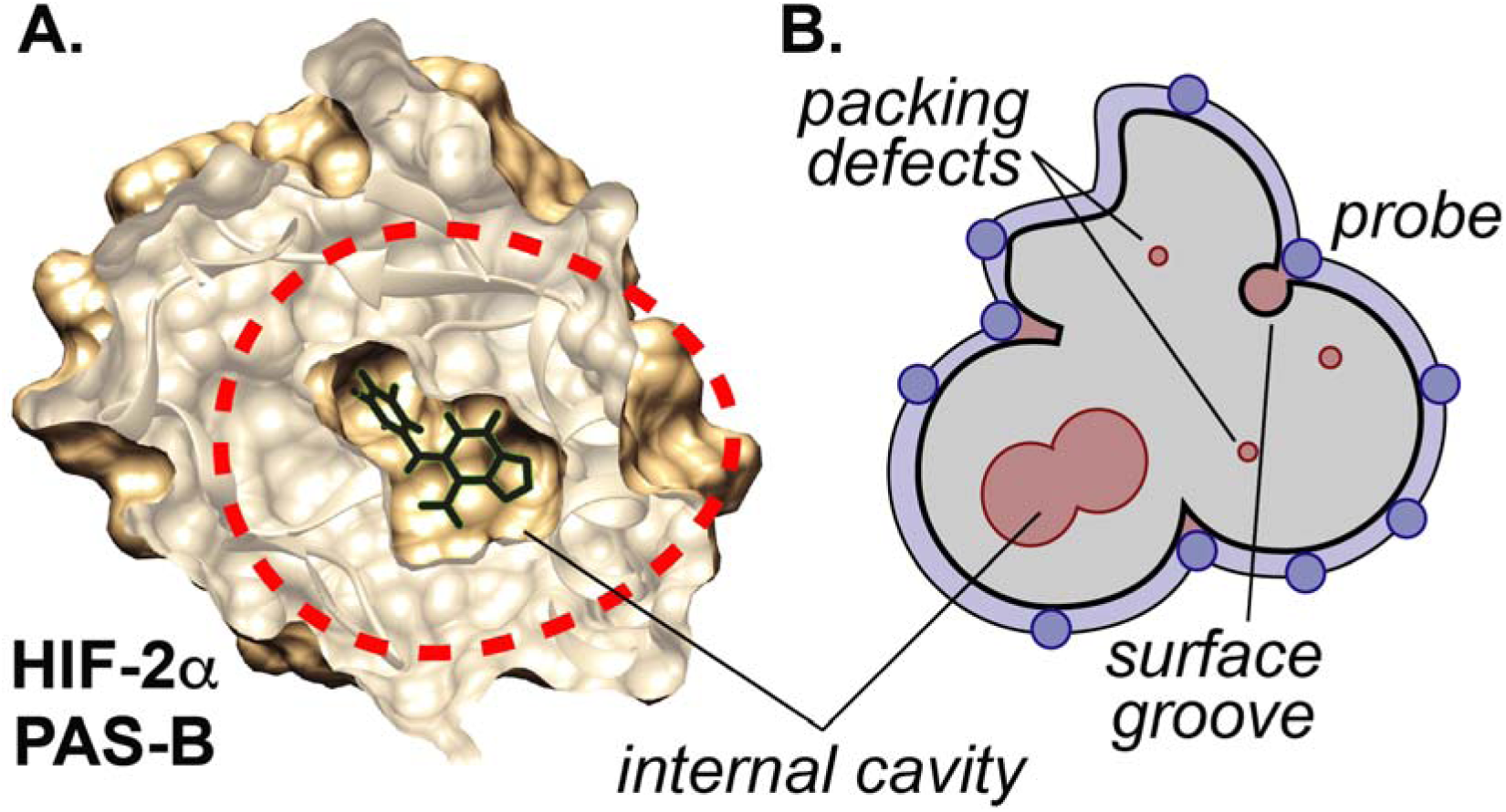
Example of internal ligand binding cavity and computational analysis. **A.** Example of protein/ligand complex utilizing an internal cavity (HIF-2*α* PAS-B, PDB: 3F1O (6)) that is sequestered over 6 Å from solvent. In the apo-form of this protein (not shown), a pre-formed 290 Å^3^ cavity with 8 crystallographically-ordered water molecules is present at this site. **B.** Schematic definitions of cavities, voids, and algorithm used by *ProteinVolume* (5) as used for Fig. 3 and onwards in the manuscript. We refer to *cavities* as internal openings larger than a single water molecule (V > 30 Å^3^ (27)), while *voids* are more generally non-protein filled spaces that include cavities along with other types of packing defects distributed throughout and around a protein. With a protein structure available (gray), the *<u>total void volume</u>* can be straightforwardly calculated by the difference of the solvent-excluded volume of a protein (generated by rolling probes over the molecular surface, typically with water-sized radii) and the volume taken by protein atoms = volume under blue surface minus gray volume (5).

The traditional method to identify such locations – high-resolution X-ray crystallography, ideally with better than 2 Å resolution to aid the particularly when combined with computational analyses for cavity identification (5, 14–21) identification of internally bound waters – can be powerful, or experimentally solving multiple structures of proteins soaked with different organic solvents or small molecule fragments (22, 23). Accordingly, this approach relies on having well-diffracting crystals, which are not always available for all systems and can be time-consuming to produce even when successful. In parallel, several classes of computational tools for predicting protein cavities and other ligand-binding sites have rapidly advanced through a variety of approaches (9, 21, 24–26). While these usefully expand the hypothesis space, they also highlight the need for fast experimental validation.

As an alternative method to experimentally determine which proteins might contain internal cavities suitable for ligand binding, we explored the potential for using high pressure solution NMR to do so. We thought this approach might be useful, given the need for proteins to undergo structural changes to allow for ligand binding within their pre-existing internal cavities and voids (**Fig. 1**). While such changes may occurrarely at ambient pressure, elevated pressure is known to dramatically increase the equilibrium populations of the less populated, low-volume conformers associated with hydration of cavities and voids (28). Such low-lying excited state conformers (N’), if present, usually equilibrate with the ground state folded conformer (N) rapidly on the NMR time scale (*τ* < ms). This gives rise to population-weighted single peaks in multidimensional spectra which exhibit *non-linear chemical shift changes* as pressure pushes the equilibrium N ⇌ N’ toward N’ (29). We anticipated that this effect might be easily detected in other globular proteins at pressures in the 500-2000 bar range, below the pressures which typically partially or completely unfold proteins (29–31).

To evaluate this approach, we examined the effects of high hydrostatic pressure on the NMR chemical shifts of a collection of protein domains and protein-ligand complexes. Many of these proteins are members of the Per-ARNT-Sim (PAS) family of ligand-controlled protein/protein interaction domains, which often internally bind different small molecule cofactors to sensitize them to environmental factors like O_2_, light, and xenobiotics (32, 33). Changes to the occupancies or configurations of these cofactors trigger conformational changes in the surrounding protein, regulating the activity of various effector domains in natural and engineered proteins (33, 34). While a number of high-resolution structures of apo- and ligand-bound PAS domains have been solved, these models and predictions which can be made from them can provide insights into only a small fraction of the many thousands of “orphan” PAS domains without known ligands. Complementing these proteins, we added additional proteins and protein/ligand complexes from a wide range of domain types – including some which are known to bind ligands internally, some not – to establish the generality of this approach for quickly probing protein structure and function.

Here we test the ability of high-pressure NMR to rapidly identify void-containing proteins, with three key advances. First, by analyzing pressure titration data from over 40 proteins and protein/ligand complexes, we show that an easily accessible metric – how differently sites within a protein respond to increasing pressure, as assessed by the diversity of non-linear chemical shift perturbations observed in a simple titration without requiring site-specific assignments – correlates well with the void volume within a protein. We find that this metric is robust enough to predict total void volume on its own, allowing the prioritization of potential ligand-binding capability amongst several targets. Second, we demonstrate that internal ligand binding within a protein can reduce this heterogeneity, quickly providing information on ligand binding and, in certain cases, mode of action. Finally, we illustrate how this method can also be used to rapidly assess the impact of point mutations/repacking, such as those used to fill or generate cavities to facilitate artificial control, on the prevention or enabling of water entry. Taken together, our data show the general utility of these rapidly acquired, easily analyzed data to provide this important biophysical characterization of new proteins.

## Results

### Evaluation of non-linear NMR chemical shift responses to high pressure

We began by examining the pressure dependence of NMR signals from a set of well-characterized proteins using the workflow in **Fig. 2**. For each U-^15^N labeled protein sample, we acquired ^1^H/^15^N HSQC spectra at increasingly higher pressures from 20-2500 bar. After each individual dataset was acquired at high pressure, we lowered the pressure to 20 bar and acquired a spectrum to confirm the reversibility of conformational changes; after any sign of irreversible changes in peak intensity or location, the pressure series was stopped. These series were typically composed of 21 spectra each taking 60 min apiece, for a total of approximately 21 hr. Post-processing, peaks were picked and the pressure dependence of changes of their ^1^H and ^15^N chemical shifts were independently fit to a second-order polynomial equation:

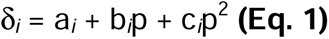

**Fig. 2:**
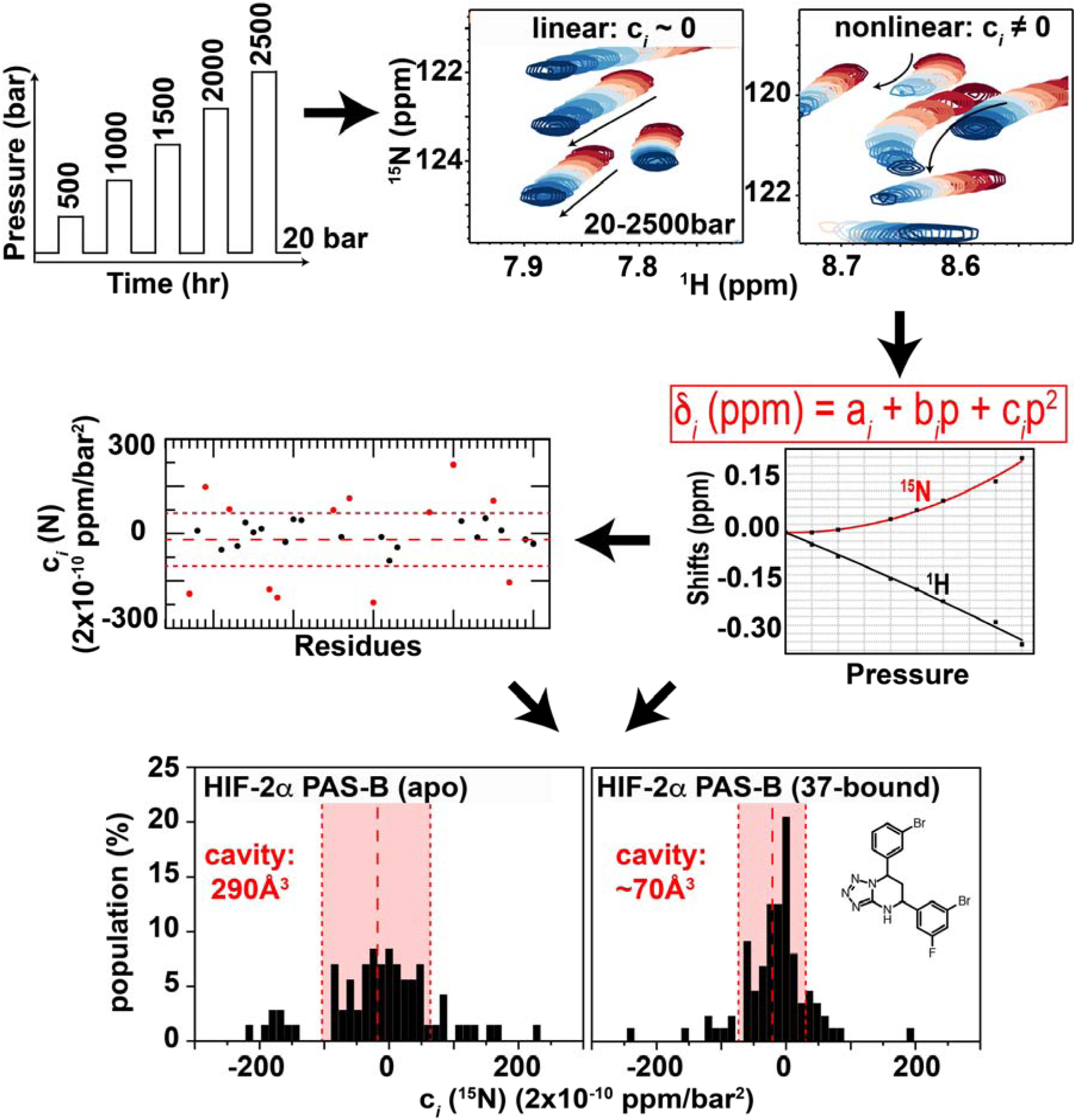
Workflow of pressure NMR analyses. For each protein analyzed, ^1^H/^15^N HSQC spectra were acquired at increasing pressures from 20-2500 bar, interleaving additional spectra at 20 bar between those at higher pressure to assess protein reversibility. Following processing, peaks were picked and chemical shifts monitored as a function of increasing pressure. Independently treating ^1^H and ^15^N movements, the pressure-dependent chemical shift changes of each peak are fit to the second order polynomial indicated to obtain linear (b*_i_*) and nonlinear (c*_i_*) coefficients. Subsequent analyses of the nonlinear (c*_i_*) coefficients include either plotting c*_i_*values as a function of residue number (for proteins with backbone chemical shift assignments) to identify regions with likely pressure-dependent conformational changes or to simply generate histograms of c*_i_* values to give a quick initial characterization of likely ability to adopt multiple folded conformations.

As established by Akasaka and co-workers (29, 30), the linear (b*_i_*) and nonlinear (c*_i_*) coefficients of these pressure responses reflect different properties of each protein. To evaluate various analyses of these data, we examined an initial group of nine proteins (“test set”) with high-resolution structures with total void volumes ranging from approximately 1500-8500 Å^3^ as assessed by ProteinVolume (5). This approach provides a sum of the volumes of a wide array of packing defects, cavities, etc. regardless of their size or distribution within a protein structure by simply calculating the difference between the solvent-accessible volume of a protein and the volume occupied by protein atoms (**Fig. 1B**). Pressure titration data were additionally recorded from more than 30 additional proteins and protein-ligand complexes, with varying degrees of structural information to build the “complete set” of data for subsequent analyses (a complete list of all analyzed proteins and complexes is provided in **Table S1**).

As an initial analysis, we examined the absolute values of the two chemical shift coefficients (|b*_i_*|, |c*_i_*|), separately averaged over all backbone amide protons and nitrogens (typically 25-125) within each protein. From these analyses, we confirmed that the averaged ^1^H and ^15^N |b*_i_*| values were fairly uniform across proteins with less than two-fold variation, while the corresponding |c*_i_*| values varied over 4-10 fold ranges (**Fig. S1**). These data expand the original data sets of Akasaka and Li (30) from seven proteins and complexes to over 40 total, add an independent measurement and analysis of the GB1 protein to assess the impact of different equipment, labs, and software for data acquisition and analyses (GB1: 750 MHz in Kobe University by K.A.; GB1’ 700-800 MHz in New York by K.H.G.), and support the original interpretations that the linear |b*_i_*| component is relatively fixed among systems while |c*_i_*| depends on a protein-specific feature (30).

Further investigating these pressure-dependent chemical shift changes, we noted a trend towards proteins with larger total void volumes having both higher c*_i_* values (**Fig. S1**) and a greater range of individual residue-specific c*_i_*parameters than proteins with smaller voids, both of which we thought might reflect a more heterogenous structural response. While the larger c*_i_* values are captured by the previously-described average |c*_i_*| parameter (30), we examined several ways to quantitate the heterogeneity in c*_i_* values, settling on a combination of histograms and the standard deviation of the c*_i_* (stdev(c*_i_*)). While the correlation between average |c*_i_*| and stdev(c*_i_*) values for amide ^1^H and ^15^N shifts (**Fig. S2**) suggest similarities between the two metrics, we opted to proceed using stdev(c*_i_*) to take advantage of the larger range of values provided by dropping the absolute value operation. We also observed a high correlation between the stdev(c*_i_*) values of ^1^H and ^15^N (**Fig. S3**), suggesting that non-linear shift changes of either nucleus report on pressure-induced changes despite differences in the structural factors which influence them (35). Our subsequent analyses utilized stdev(c*_i_*) of ^15^N chemical shift changes (= stdev(c*_i_* [^15^N])), which are thought to be most strongly influenced by changes in backbone torsion angles (30, 35).

To examine the linkage of stdev(c*_i_*) to total void volume, we measured the nonlinear components (c*_i_* [^15^N]) of a test set of nine proteins (with one, GB1, duplicated as noted above) with known structures and a range of total void volumes (**Fig. 3**).

**Fig. 3:**
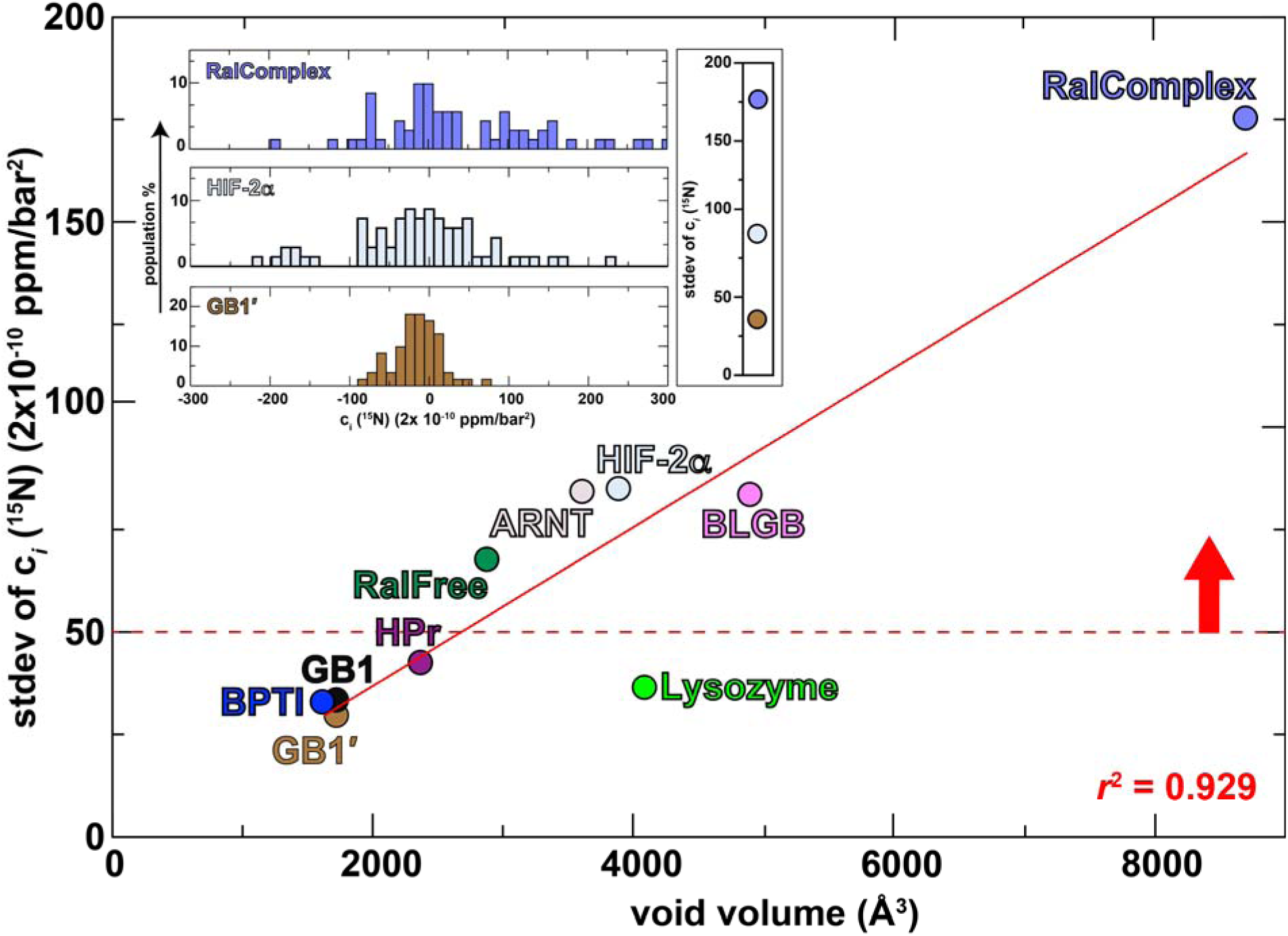
Increasing diversity of non-linear pressure dependent chemical shift changes correlates with increased void volume. Total void volumes calculated by ProteinVolume (5) for each protein are presented on the x-axis; the y-axis plots the standard deviation of the nonlinear c*i* parameters for the amide 15N chemical shifts, using data from 25-125 crosspeaks for each protein. Note that the GB1 protein was analyzed from independently produced and acquired data from two labs, as noted in the text. The linear regression (red line, y= – 1.564+0.0194x, omitting the lysozyme point) shows a correlation coefficient (*r*2) of 0.929. The red dashed line indicates an arbitrary value of approximately 50 (x2×10-10) ppm/bar2 as a recommended minimum value for potential small molecule binding cavities, as noted in the text. Inset: histograms of measured 15N c*i* parameters for three selected proteins (using the same color scheme as main figure; histograms for all ten proteins provided in **Fig. S8**).

Histograms of stdev(c*_i_*) parameters exhibited the previously mentioned variability, with proteins having larger total void volumes typically having broader distributions than those with smaller total void volumes (**Figs. 3 inset** and **S8**). We interpreted this trend as stemming from voids enabling proteins to increasingly shift from the native N conformation to a second folded N’ conformation under pressure, reflected in the non-linear chemical shift responses as the N’ is progressively populated. Quantitating the breadth of these distributions by stdev(c*_i_* [^15^N]) and plotting these versus total void volume showed a linear correlation coefficient (*r*^2^) of 0.929, supporting the linkage between pressure-induced chemical shift nonlinearities to identify cavities. Of note, the proteins with a stdev(c*_i_*[^15^N]) above 50 (x 2×10^-10^ ppm/bar^2^) are all known to bind other proteins or small molecules, suggesting an arbitrary value which could be useful to prioritize for ligand-screening.

We note that the primary outlier of the observed linear correlation, hen egg white lysozyme, is thermostable with a *T*_m_ of almost 75°C (36), likely hampering its transition into an excited state under pressure unless its intrinsic stability of its basic folded state is substantially lowered (*e.g.* by cooling close to the cold denaturation temperature (37)). In addition, we cannot rule out potential contributions from lysozyme being an enzyme instead of a signal transduction component, which may contribute to a different ability to adopt alternative conformations reflected in pressure-induced non-linear chemical shift changes. This remains to be explored more broadly in future studies.

### Use of pressure NMR to reveal ligand binding mode of action – Well-defined cases

Our correlation between increased heterogeneity of pressure-induced chemical shift changes and void size makes a strong prediction that this route should provide a rapid way to assess ligand binding: ligands which bind within the protein and reduce total void volume should decrease the stdev(c*_i_*) value of spectra recorded on the receptor. To test this prediction, we evaluated how ligand binding affects the PAS-B domains of the human HIF-2*α* and ARNT proteins, both of which are involved in the human hypoxia response (38) and contain internal cavities known to bind artificial small molecule ligands (1, 3, 6).

Prior studies of the HIF-2*α* PAS-B domain from us and others (1, 6, 39, 40) have shown that it contains an internal water-filled 290 Å^3^ cavity with no obvious access to external solvent. However, a variety of screening efforts – from NMR-based fragment binding screens to high throughput screens of HIF-2*α*/ARNT disruption (1, 2, 6, 40) or protein stability (40) – have identified a wide range of small molecules which bind into this cavity with nano- to micromolar affinities. High-resolution structures of these complexes show that the ligands displace water and reduce HIF-2 activity by impacting HIF-2*α*/ARNT interactions (41–43). To accommodate these binding events within a solvent-inaccessible cavity, the HIF-2*α* PAS-B protein must dynamically fluctuate to allow the entry of small molecules into the interior (44). Our pressure NMR analysis confirmed this hypothesis, showing a very broad distribution of c*_i_* (^15^N) responses with a correspondingly large stdev(c*_i_* [^15^N]) of 85 x (2×10^-10^ ppm/bar^2^) (**Figs. 4A, S4**). We anticipated that repeating these measurements in the presence of two nanomolar-affinity compounds (2 (1) and 37 (2)) would show smaller non-linear chemical shift changes than the apo protein given the smaller total void volume and expected reduced flexibility of the protein/ligand complexes. Our data supported this, as we observed decreases in stdev(c*_i_* [^15^N]) from 85 x (2×10^-10^ ppm/bar^2^) for the apo protein to 64 (2-bound) and 53 (37-bound) x (2×10^-10^ ppm/bar^2^), respectively. Notably, these decreases do not solely correlate with loss of void volume in some cases: measurements conducted on HIF-2*α* D1, a computationally-repacked variant with five point mutations which reduce the cavity volume to 77 Å^3^ (45) while retaining function, shows an *increase* in stdev(c*_i_* [^15^N]) up to 131 x (2×10^-10^ ppm/bar^2^) (**Fig. 4A**). We speculate that some dynamic or thermodynamic stability aspect of this engineered protein differs markedly natural proteins, contribute to its unusual pressure response.

**Fig. 4:**
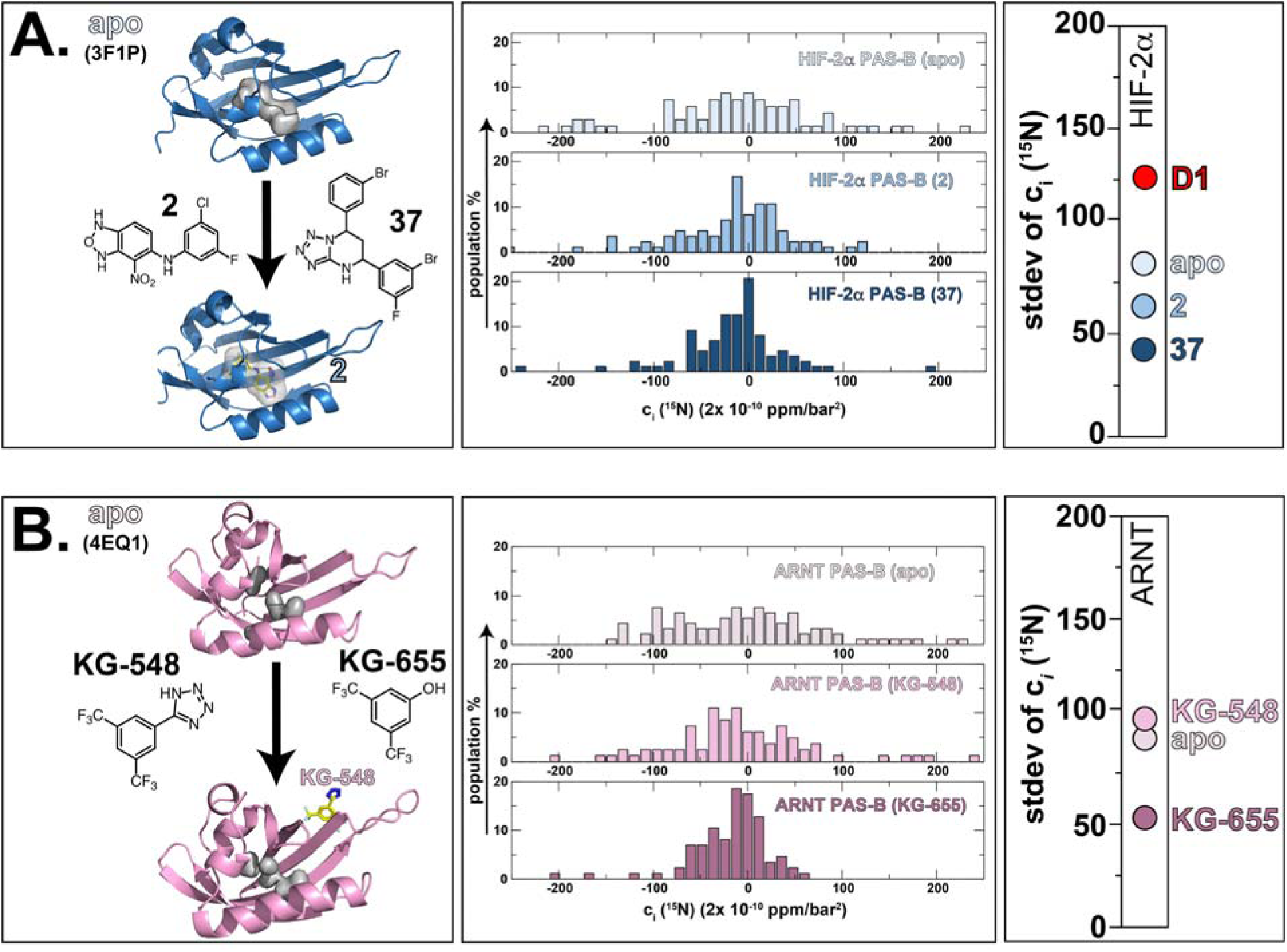
High pressure NMR can provide mode-of-action information on ligand binding, even without structural information. (Left panel) **A**. Ribbon diagrams of HIF-2*α* PAS-B in its apo (PDB: 3F1P) and holo forms (complexed with 2, PDB: 4GHI), highlighting the location of the internal ligand binding cavity. Structures of two high-affinity inhibitors, 2 and 37 (1, 2) are also shown. **B**. Ribbon diagram of ARNT PAS-B (apo PDB: 4EQ1; KG-548 complex PDB: 8G4A) with the location of internal cavities, along with the structures of two moderate affinity binders, KG-548 and KG-655 (3). No structural details were available for a compound-bound form at the outset of this work. (Central panel) Histograms of the ^15^N c*_i_* coefficients measured on apo- and holo forms of HIF-2*α* and ARNT PAS-B domains using the approach diagrammed in Fig. 2. (Right panel) Comparisons of the stdev(c*_i_* [^15^N]) values of HIF-2*α* and ARNT PAS-B apo and ligand-bound samples, showing substantial effects upon binding small molecules (or point mutations, in the case of HIF-2*α* PAS-B “D1” variant) into the internal cavities.

We used the same approach with the ARNT PAS-B domain, which has the same fold as HIF-2*α* PAS-B but contains smaller internal cavities totaling 150 Å^3^ in volume. We have previously used NMR-based fragment screening, isothermal titration calorimetry (ITC) and microscale thermophoresis (MST) to identify several compounds which bind ARNT PAS-B with micromolar affinities; two of these, KG-548 and KG-655, are further known to disrupt ARNT PAS-B interactions with coactivator proteins (3). However, the binding modes of these compounds remained unclear without NMR or X-ray co-complex structures available at the outset of this work.

To gain more insight on how KG-548 and KG-655 bound ARNT PAS-B, we used our pressure NMR analysis, confirming that ARNT PAS-B retains a flexibly-accessible interior cavity with a stdev(c*_i_* [^15^N]) of 84 x (2×10^-10^ ppm/bar^2^) (**Figs. 4B, S5**). Repeating these in the presence of the KG-655 ligand, we observed a drop of a stdev(c*_i_*[^15^N]) to 51 x (2×10^-10^ ppm/bar^2^), consistent with an interior binding location for this small ligand analogous to the HIF-2*α* PAS-B examples above. However, similar studies with the larger KG-548 ligand show a stdev(c*_i_* [^15^N]) comparable to the apo protein, with an observed value of 96 x (2×10^-10^ ppm/bar^2^) that suggests that this compound binds in a different mode than KG-655, potentially outside the cavity. We subsequently verified this external binding mode for KG-548 by determining the X-ray crystal structure of ARNT PAS-B with this ligand soaked into the crystal (**Table S2**). The resulting 1.97 Å model (**Fig. S9**) clearly demonstrated that KG-548 bound across the outside of the PAS*β*-sheet, which we independently validated in solution (46).

### Use of pressure NMR to reveal ligand binding – Poorly-defined cases

As a next demonstration of this approach, we examined its utility for PAS domains outside the hypoxia response and with less well understood regulation. We considered two such targets, the first of which is the N-terminal PAS domain of human PAS kinase (hPASK PAS-A), a PAS domain-regulated serine/threonine kinase conserved among eukaryotes (4, 47). This domain has a canonical PAS structure, with a five-stranded *β*-sheet flanked to one side by several *α* helices (32, 48). While only a small surface groove of 73 Å^3^ could be identified near the F/G loop in the representative member of the solution structure ensemble, an NMR-based fragment screen of 750 compounds identified several small molecules that bound with micromolar affinity to hPASK PAS-A (4). Chemical shift perturbations suggest that these compounds bind within the domain interior, analogous to the subsequently discovered HIF-2*α* and ARNT PAS-B binding compounds, but the lack of a hPASK PAS-A/ligand complex structure leaves this as an open issue. To address this, we used high-pressure NMR to examine the response of hPASK PAS-A in its apo form and when saturated with two compounds with *K*_D_=13-24 µM affinities (KG-535, KG-571 = compounds 1 and 2 in (4)). For the apo-protein, we observed a stdev(c*_i_* [^15^N]) of 45 x (2×10^-10^ ppm/bar^2^); from the correlation identified in our test set (**Fig. 3**), this value predicts that hPASK PAS-A contains a total void volume slightly smaller than the 3345 Å^3^ calculated from the representative member of the solution structure ensemble (**Figs. 5A, S6**). Upon addition of the KG-535 and KG-571 ligands, we observed smaller stdev(c*_i_* [^15^N]) values for both ligand-bound forms (35 and 28 x (2×10^-10^ ppm/bar^2^) for the KG-535 and KG-571-bound forms, respectively (**Fig. 5A**). These data suggest that both compounds bind within the void volume of hPASK PAS-A, with KG-571 having a greater effect on the domain adopting alternative folded conformations. Additionally, Boltz-2 (7) predicted holo structures of hPASK PAS-A shows both ligands binding within an internal cavity (**Figs. 5B, S10B**), consistent with our pressure NMR findings.

**Fig. 5:**
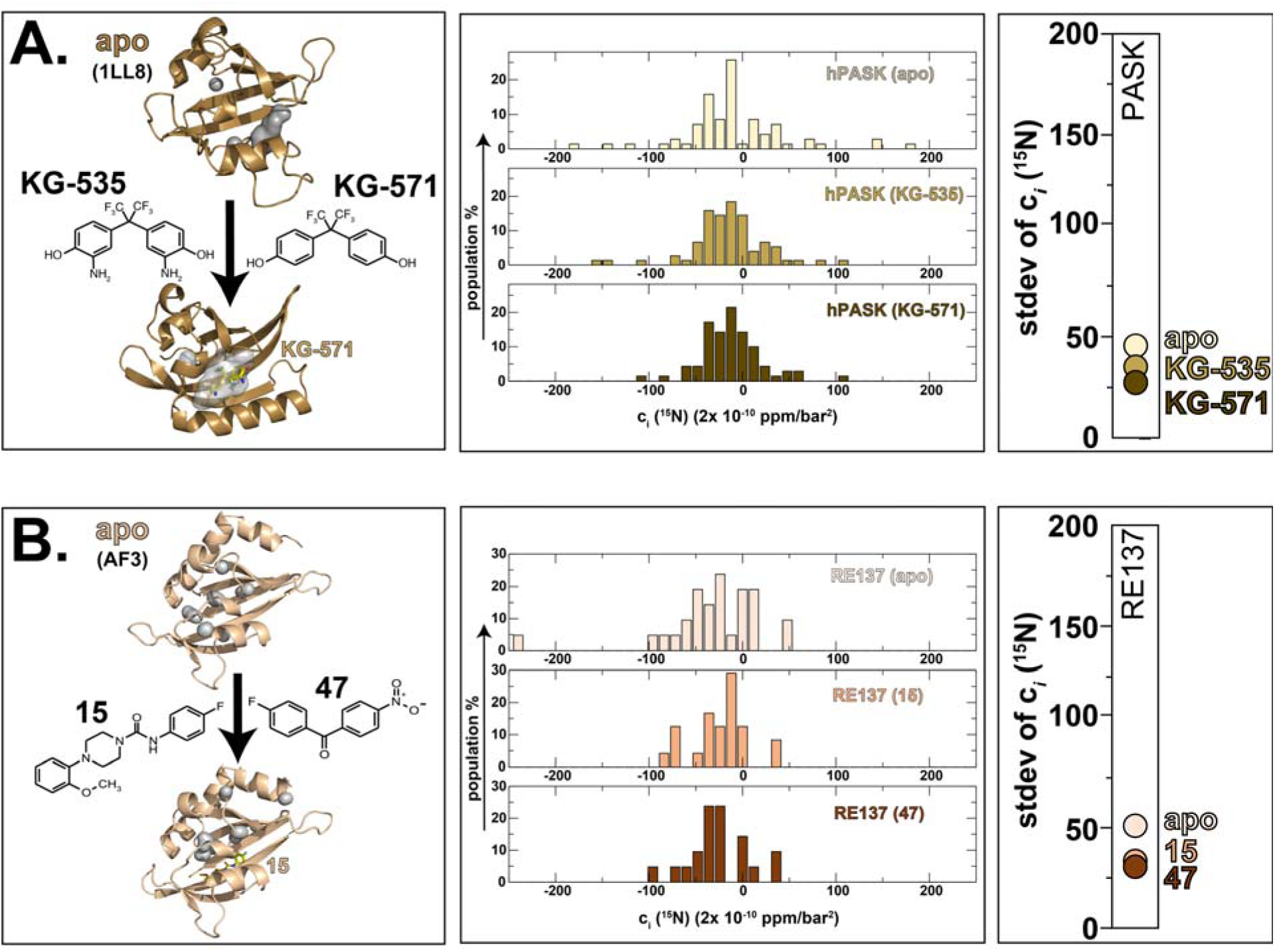
High pressure NMR can provide ligand binding information even in the absence of experimental structure. (Left panel) **A.** Three-dimensional structure of hPASK PAS-A (PDB: 1LL8) and structures of two binding compounds, KG-535 and KG-571 (4). No high-resolution structures of protein/ligand complexes are available, protein/ligand complex was generated using Boltz-2 (7). **B.** No experimental structural information is available for the RE137 PAS domain (8) (either apo or bound forms), structure shown here was generated using AlphaFold3 (AF3) (9, 10). Structures of two binding compounds identified by ^19^F ligand-detected NMR screening, 15 and 47 and the protein/ligand complex was generated using Boltz-2 (7). (Central panel) Histograms of the ^15^N c*_i_* coefficients measured on apo- and holo forms of hPASK and RE137 PAS domains using the approach diagrammed in Fig. 2. (Right panel) Comparisons of the stdev(c*_i_* [^15^N]) values of hPASK and RE137 apo- and ligand-bound samples.

We completed our analyses of natural proteins by examining a prokaryotic PAS domain from a novel histidine kinase from *Rhizobium etli* (*8, 49*), chosen because of a). homology with the light-regulated PAS-histidine kinase EL346 (50, 51) and b). the widespread usage of small molecule ligands to control the structure and function of PAS-containing proteins (including many histidine kinases (33)). While no experimental structures are available of this domain, which we designate RE137, we hypothesized that it may contain a cavity that can accommodate a small molecule, given precedence from other PAS domains (33) and the presence of several small cavities within an AlphaFold3 (AF3) (9) model of apo RE137 (**Fig. S10B**). Pressure NMR analyses gave experimental support for this hypothesis, as the stdev(c*_i_* [^15^N]) value of 53 x (2×10^-10^ ppm/bar^2^) (**Figs. 5B, S7**) we obtained from the apo protein is consistent with a 2800 Å^3^ total void volume from the correlation seen in **Fig. 3**. To provide an initial assessment of small molecule binding to RE137, we used an *R*_2_-filtered ^19^F NMR assay to screen a 58-member library of fluorinated compounds (52) for their binding to this protein. Two compounds from this screen showed binding, 15 and 47; subsequent pressure NMR analyses of RE137 in presence of 1.2 mM of either compound exhibited decreases of 43% and 39% with stdev(c*_i_* [^15^N]) values of 30 and 32 x (2×10^-10^ ppm/bar^2^) for the 15-and 47-bound forms, respectively. We interpret these data to indicate that RE137 can bind both compounds in a manner that reduces the total void volume of the protein, consistent with interior binding locations for these small ligands.

### Use of pressure NMR to probe artificially-designed proteins

As a final demonstration of the utility of this method, we examined its utility in investigating the flexibility of an artificially-designed protein, CA01 (53). With advances in protein engineering enabling the development of artificial ligand-binding biosensor proteins (54), we sought to probe whether a protein such as CA01 – which contains a completely-enclosed 75 Å^3^ cavity within approximately 3500 Å^3^ of total void volume – might show similar pressure-dependent non-linear chemical shift changes as natively-evolved counterparts. From our correlation in **Fig. 3** we expected to see a stdev(c*_i_* [^15^N]) value of approximately 75 (2×10^-10^ ppm/bar^2^) from a natural protein with void volume of this size, but we instead observed a substantially smaller value of 38 x (2×10^-10^ ppm/bar^2^) (**Fig. 6**). As CA01 is extremely stable to thermal and chemical denaturation, requiring 5 M guanidinium hydrochloride to exhibit a complete thermal melt with a 75°C *T*_m_ (53), we view this observation supports our prior findings with lysozyme suggesting thermostable proteins being less able to adopt alternative conformations (**Fig. 3**).

**Fig. 6:**
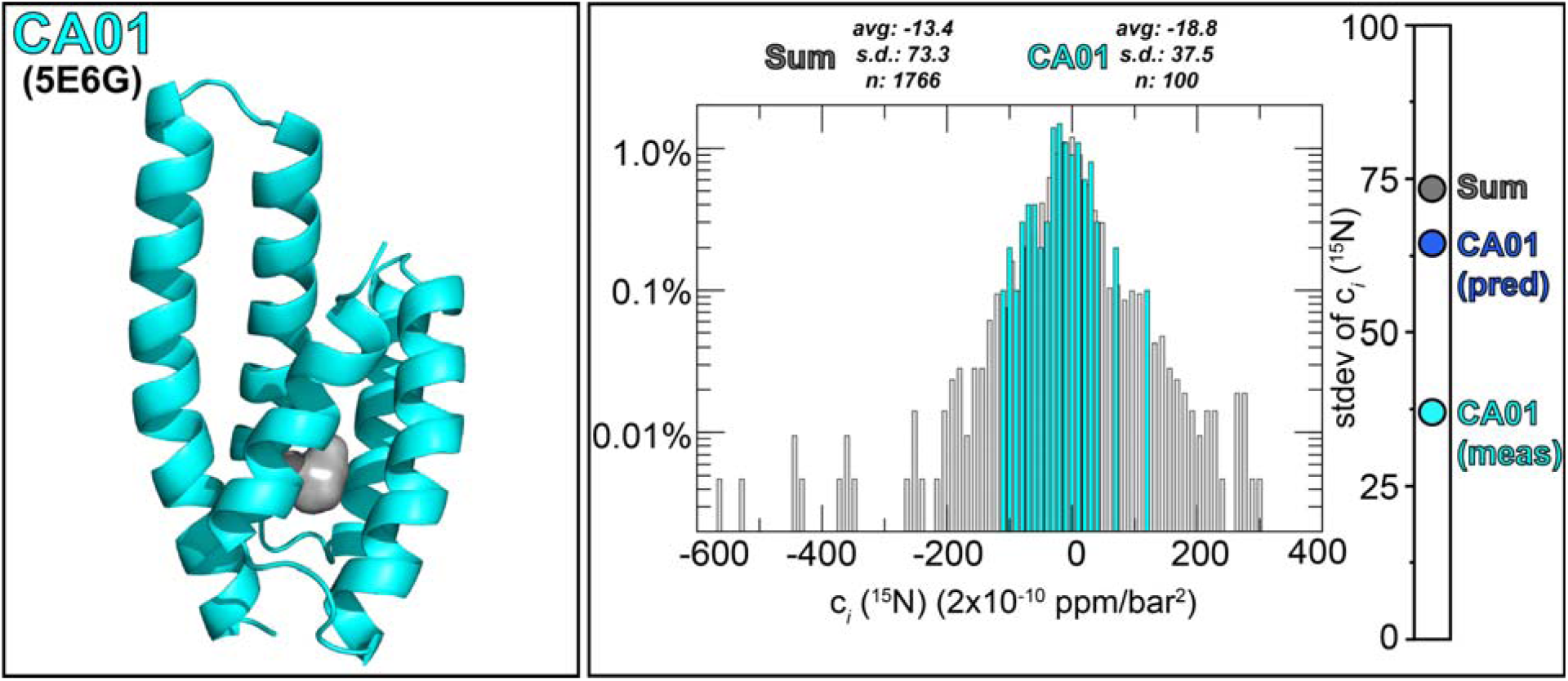
High pressure NMR of a thermostable designed protein with an internal cavity gives smaller non-linear pressure dependent chemical shift changes than anticipated from void volume. (Left panel) Three-dimensional structure of CA01 (PDB: 5E6G), highlighting the location of a 75 Å3 internal cavity within a total void volume of 3500 Å3. (Right panel) Histogram of the 15N c*i* coefficients measured on CA01 (aqua) and 22 apo proteins listed in **Fig. S1** (grey) using the approach diagrammed in Fig. 2. Bar on right indicates stdev(c*i* [15N]) measured for CA01 (aqua) and 22 apo proteins (grey), along with the value predicted for CA01 from the correlation in Fig. 3 (blue).

## Discussion

In the present work, we successfully used non-linear pressure-induced chemical shift effects at backbone amide ^1^H and ^15^N nuclei to rapidly assess whether proteins contain substantial total void volumes within them and if these can bind small molecule ligands. A particular strength of our implementation is its simplicity, as 1). U-^15^N labeled samples are routinely available from *E. coli* or *in vitro* expression systems (or otherwise, using natural abundance ^15^N samples with optimized data collection), 2). NMR data needed are easily obtained from conventional ^1^H/^15^N HSQC experiments recorded at variable pressures using commercially available pressure equipment, and 3). straightforward data analyses are done by simply tracking peak locations without requiring chemical shift assignments, eliminating a time-consuming step of NMR analysis. These characteristics make our method broadly accessible, data-rich, and fast enough to integrate with emerging discovery pipelines.

Mechanistically, we attribute the non-linear chemical shift changes with increasing pressure to site-specific conformational changes of the protein from the native folded structural ensemble (N) to the low-lying excited state ensemble (N’) with a smaller volume (17). Our specific choice of backbone amide ^1^H and ^15^N chemical shifts give us complementary views of these transitions (**Fig. S3**), as ^1^H chemical shifts are dominated by hydrogen bonding to these atoms, while ^15^N chemical shifts are influenced by a broader set of factors, including changes in hydrogen bonding (to both the adjacent ^1^H and carbonyl CO groups), local backbone dihedrals and sidechain conformations (55). Of note, non-linear pressure-dependent chemical shift changes at these sites are much smaller in peptides or intrinsically disordered proteins (56). Indeed, non-linear pressure-dependent amide ^1^H and ^15^N chemical shift changes have long been recognized as reflecting idiosyncratic differences among proteins (57). Prior work by one of our groups (K.A.) suggested that this property depends on the density of cavities within the protein (30); here we both expand the number of proteins examined from that work and suggest that contributions likely arise from small and large voids distributed throughout the protein which collectively give rise to the total void volumes calculated by the program ProteinVolume (5) given the excellent correlation we present in **Fig. 3**. We posit that such an approach reveals aspects that will be correlated to the ability of proteins to be similarly affected by allosteric regulators and other ligands, linking these non-linear chemical shift effects to the potential for small molecule binding and regulation.

In addition to getting information on the total void volume within a given protein, the strengths of this analysis include the ability to identify ligands which bind to a given target and (de)stabilize one conformation in a way that may provide allosteric switching. As such, this provides trivially accessible mode-of-action information that is often challenging to get otherwise without resources such as compounds of known binding location for competition studies or cavity-filling point mutants. Our comparison of two cavity-perturbing effects for HIF-2*α* PAS-B – small molecule binding and cavity-filling redesign – produced opposite effects, with the small molecules stabilizing the native fold and mutations destabilizing in our pressure NMR analyses. We underscore that these effects occur despite minor changes in ground state structure among structures of the apo, ligand-bound and redesigned proteins (1, 6, 45). As such, pressure-dependent chemical shift effects give us insight into details of the packing and dynamics of proteins more simply than most other methods can provide. Finally, our comparison of the ARNT PAS-B binding compounds KG-548 and KG-655 provided an outstanding example of the power of the pressure NMR method to quickly and accurately provide mode-of-action binding site information even with limited structural data available. In these cases, high pressure NMR – using rapidly-acquired spectra without needing any chemical shift assignments – indicated different binding modes between the two ligands, with KG-655 binding primarily to an internal site while KG-548 bound externally. These findings were subsequently validated both by a ARNT PAS-B:KG-548 crystal structure (**Fig. S9**) and accompanying solution biophysical investigation (46).

We emphasize that our approach requires very little preliminary information on the target, needing neither structures nor site-specific chemical shift assignments, as demonstrated by the RE137 case. In this case, the stdev(c*_i_* [^15^N]) values of the apo protein exceeds the value of approximately 50 x (2×10^-10^ ppm/bar^2^) value that we suggest as indicating the potential for ligand binding, and we were able to quickly identify compounds which bound RE137. Our high pressure NMR results quickly add important information, showing that these compounds bind and modulate various aspects of protein structure and dynamics, probably through binding internal cavities.

A final strength of the method is its speed, making this pressure NMR method an outstanding complementary experimental tool to computational pipelines which can predict protein cavities (21) or protein ligand interactions (9, 24–26) at scale. While powerful, these predictions require experimental validation to confirm which proposed interactions translate into meaningful binding events in solution. Via ligand-induced changes in stdev(c*_i_* [^15^N]) values, pressure NMR provides a rapid experimental screen to identify proteins and small molecule binders worth pursuing with higher resolution experimental NMR, X-ray, or cryo-EM analyses.

Finally, while our demonstrations have focused on relatively small ∼15 kDa ligand binding domains, we note that several routes can expand the reach of this approach to larger and more complex systems by reducing spectral complexity. Options include biochemical approaches to generate proteins with simplified ^15^N-labeling patterns, as provided by amino-acid type selective or sortase/intein-based domain-specific techniques, to ^1^H/^15^N TROSY or HNCO-type pulse sequences which can provide improved resolution. Additional options may be provided by the improved NMR signal properties of ^1^H/^13^C methyl signals for similar pressure-sensitive studies, particularly given the placement of these residues near internal cavities.

## Materials and Methods

Proteins were expressed in *E. coli* with uniform ^15^N labeling, purified using a combination of Ni(II) affinity and gel filtration chromatography before being exchanged into a barostatic Tris:phosphate buffer mix which limits pH changes during pressurization (58) and concentrated to 100-700 µM for NMR spectroscopy. ^1^H/^15^N HSQC spectra were acquired at increasing pressures from 20-2500 bar, interleaving each high pressure spectrum with a low pressure (20 bar) spectrum to confirm that changes in peak locations and intensities were reversible. All NMR data were processed and analyzed with NMRviewJ (59, 60) and NMRFx (59, 61, 62). After individually processing each spectrum, we picked peaks and tracked their changes in chemical shifts as a function of pressure. Separately handling movement in the ^1^H and ^15^N dimensions, we fit these trends to a second-order polynomial equation (**Eq. 1**, **Fig. 2**). Protein volume analyses of single isolated cavities utilized *cavfinder* (1), while measurements of total void volumes utilized *ProteinVolume* (5). Additional detailed procedures are found in **SI Materials and Methods**.

## Acknowledgments

The authors gratefully acknowledge assistance in sample preparation from Dong Lee and Andrew Palacios (CUNY ASRC), and comments on the manuscript by Hua Li (Chinese Academy of Sciences, Shanghai). U-^15^N samples of FABP proteins were generously provided by May Poh Lai and Ruth Stark (CCNY); plasmids of other proteins were provided by Elena Rusinova and Ming-Ming Zhou (Mt. Sinai Medical Center, BRD proteins), and T.M. Jacobs and Brian Kuhlman (UNC Chapel Hill, CA01). This work was supported by NIH grants R01 GM106239 (K.H.G.), R35 GM156296 (K.H.G.), T34 GM007639 (supporting L.P.M.), F31 GM142258 (R.A.), R01 GM123012 (B.A.J.), and U54 AI170660 (B.A.J.), NSF grant MCB-1818148 (K.H.G.), as well as a postdoctoral fellowship to D.G. (number B3X from the Fonds de Recherche Québec – Nature et Technologie (FRQNT)).

This work is based upon research conducted at the Northeastern Collaborative Access Team beamlines, which are funded by the National Institute of General Medical Sciences from the National Institutes of Health (P30 GM124165). The Eiger 16M detector on the 24-ID-E beam line is funded by a NIH-ORIP HEI grant (S10OD021527). This research used resources of the Advanced Photon Source, a U.S. Department of Energy (DOE) Office of Science User Facility operated for the DOE Office of Science by Argonne National Laboratory under Contract No. DE-AC02-06CH11357. Additional data were acquired using the resources 17-ID-1 of the National Synchrotron Light Source II, a U.S. Department of Energy (DOE) Office of Science User Facility operated for the DOE Office of Science by Brookhaven National Laboratory under Contract No. DE-SC0012704.

## References

1. T. H. Scheuermann et al., Allosteric inhibition of hypoxia inducible factor-2 with small molecules. Nat Chem Biol 9, 271–276 (2013).

2. T. H. Scheuermann et al., Isoform-Selective and Stereoselective Inhibition of Hypoxia Inducible Factor-2. J Med Chem 58, 5930–5941 (2015).

3. Y. Guo et al., Regulating the ARNT/TACC3 axis: multiple approaches to manipulating protein/protein interactions with small molecules. ACS Chem Biol 8, 626–635 (2013).

4. C. A. Amezcua, S. M. Harper, J. Rutter, K. H. Gardner, Structure and interactions of PAS kinase N-terminal PAS domain: model for intramolecular kinase regulation. Structure 10, 1349–1361 (2002).

5. C. R. Chen, G. I. Makhatadze, ProteinVolume: calculating molecular van der Waals and void volumes in proteins. BMC Bioinformatics 16, 101 (2015).

6. T. H. Scheuermann et al., Artificial ligand binding within the HIF2alpha PAS-B domain of the HIF2 transcription factor. Proc Natl Acad Sci U S A 106, 450–455 (2009).

7. S. Passaro et al., Boltz-2: Towards Accurate and Efficient Binding Affinity Prediction. bioRxiv 10.1101/2025.06.14.659707, 2025.2006.2014.659707 (2025).

8. I. Dikiy et al., Diversity of function and higher-order structure within HWE sensor histidine kinases. J Biol Chem 299, 104934 (2023).

9. J. Abramson et al., Accurate structure prediction of biomolecular interactions with AlphaFold 3. Nature 630, 493–500 (2024).

10. R. Evans et al., Protein complex prediction with AlphaFold-Multimer. bioRxiv 10.1101/2021.10.04.463034, 2021.2010.2004.463034 (2022).

11. J. Ashkani, D. J. Rees, The Critical Role Of VP1 In Forming The Necessary Cavities For Receptor-mediated Entry Of FMDV To The Host Cell. Sci Rep 6, 27140 (2016).

12. J. Mondal, N. Ahalawat, S. Pandit, L. E. Kay, P. Vallurupalli, Atomic resolution mechanism of ligand binding to a solvent inaccessible cavity in T4 lysozyme. PLoS Comput Biol 14, e1006180 (2018).

13. A. S. Tanwar, V. D. Goyal, D. Choudhary, S. Panjikar, R. Anand, Importance of Hydrophobic Cavities in Allosteric Regulation of Formylglycinamide Synthetase: Insight from Xenon Trapping and Statistical Coupling Analysis. PLoS ONE 8, e77781 (2013).

14. C. D. Andersson, B. Y. Chen, A. Linusson, Mapping of ligand-binding cavities in proteins. Proteins: Structure, Function, and Bioinformatics 78, 1408–1422 (2010).

15. Z. Guo et al., Identification of Protein–Ligand Binding Sites by the Level-Set Variational Implicit-Solvent Approach. Journal of Chemical Theory and Computation 11, 753–765 (2015).

16. Z. Zhang, Y. Li, B. Lin, M. Schroeder, B. Huang, Identification of cavities on protein surface using multiple computational approaches for drug binding site prediction. Bioinformatics 27, 2083–2088 (2011).

17. J. Dundas et al., CASTp: computed atlas of surface topography of proteins with structural and topographical mapping of functionally annotated residues. Nucleic Acids Res 34, W116–W118 (2006).

18. J.-K. Kim et al., BetaVoid: Molecular voids via beta-complexes and Voronoi diagrams: BetaVoid: Molecular Voids. Proteins: Structure, Function, and Bioinformatics 82, 1829-1849 (2014).

19. K. P. Tan, T. B. Nguyen, S. Patel, R. Varadarajan, M. S. Madhusudhan, Depth: a web server to compute depth, cavity sizes, detect potential small-molecule ligand-binding cavities and predict the pKa of ionizable residues in proteins. Nucleic Acids Res 41, W314–W321 (2013).

20. Y. Xu et al., CavityPlus: a web server for protein cavity detection with pharmacophore modelling, allosteric site identification and covalent ligand binding ability prediction. Nucleic Acids Res 46, W374–W379 (2018).

21. A. Meller et al., Predicting locations of cryptic pockets from single protein structures using the PocketMiner graph neural network. Nat Commun 14, 1177 (2023).

22. D. A. Keedy et al., An expanded allosteric network in PTP1B by multitemperature crystallography, fragment screening, and covalent tethering. Elife 7, e36307 (2018).

23. C. Mattos, D. Ringe, Locating and characterizing binding sites on proteins. Nat Biotechnol 14, 595–599 (1996).

24. S. Passaro, et al., Boltz-2: Towards Accurate and Efficient Binding Affinity Prediction. bioRxiv 10.1101/2025.06.14.659707 (2025).

25. A. Morehead, J. Cheng, FlowDock: Geometric Flow Matching for Generative Protein-Ligand Docking and Affinity Prediction. ArXiv (2025).

26. L. Herron et al., DiffDock-Glide: a hybrid physics-based and data-driven approach to molecular docking. bioRxiv 10.1101/2025.06.02.657461, 2025.2006.2002.657461 (2025).

27. M. Gerstein, C. Chothia, Packing at the protein-water interface. Proc Natl Acad Sci U S A 93, 10167–10172 (1996).

28. K. Akasaka, Exploring the entire conformational space of proteins by high-pressure NMR. Pure and Applied Chemistry 75, 927–936 (2003).

29. K. Akasaka, Probing conformational fluctuation of proteins by pressure perturbation. Chem Rev 106, 1814–1835 (2006).

30. K. Akasaka, H. Li, Low-Lying Excited States of Proteins Revealed from Nonlinear Pressure Shifts in ^1^H and ^15^N NMR. Biochemistry 40, 8665–8671 (2001).

31. M. Gross, R. Jaenicke, Proteins under pressure. The influence of high hydrostatic pressure on structure, function and assembly of proteins and protein complexes. Eur J Biochem 221, 617–630 (1994).

32. B. L. Taylor, I. B. Zhulin, PAS domains: internal sensors of oxygen, redox potential, and light. Microbiol Mol Biol Rev 63, 479–506 (1999).

33. J. T. Henry, S. Crosson, Ligand-binding PAS domains in a genomic, cellular, and structural context. Annu Rev Microbiol 65, 261–286 (2011).

34. A. Losi, K. H. Gardner, A. Moglich, Blue-Light Receptors for Optogenetics. Chem Rev 118, 10659–10709 (2018).

35. K. Akasaka et al., Pressure response of protein backbone structure. Pressure-induced amide ^15^N chemical shifts in BPTI. Protein Sci 8, 1946–1953 (1999).

36. T. Knubovets, J. J. Osterhout, P. J. Connolly, A. M. Klibanov, Structure, thermostability, and conformational flexibility of hen egg-white lysozyme dissolved in glycerol. Proc Natl Acad Sci U S A 96, 1262–1267 (1999).

37. Y. O. Kamatari, L. J. Smith, C. M. Dobson, K. Akasaka, Cavity hydration as a gateway to unfolding: an NMR study of hen lysozyme at high pressure and low temperature. Biophys Chem 156, 24–30 (2011).

38. W. G. Kaelin, Jr., P. J. Ratcliffe, Oxygen sensing by metazoans: the central role of the HIF hydroxylase pathway. Mol Cell 30, 393–402 (2008).

39. D. Wu, N. Potluri, J. Lu, Y. Kim, F. Rastinejad, Structural integration in hypoxia-inducible factors. Nature 524, 303–308 (2015).

40. D. Wu et al., Bidirectional modulation of HIF-2 activity through chemical ligands. Nat Chem Biol 15, 367–376 (2019).

41. W. Chen et al., Targeting renal cell carcinoma with a HIF-2 antagonist. Nature 539, 112–117 (2016).

42. H. Cho et al., On-target efficacy of a HIF-2alpha antagonist in preclinical kidney cancer models. Nature 539, 107–111 (2016).

43. E. M. Wallace et al., A Small-Molecule Antagonist of HIF2alpha Is Efficacious in Preclinical Models of Renal Cell Carcinoma. Cancer Res 76, 5491–5500 (2016).

44. J. Key, T. H. Scheuermann, P. C. Anderson, V. Daggett, K. H. Gardner, Principles of ligand binding within a completely buried cavity in HIF2alpha PAS-B. J Am Chem Soc 131, 17647–17654 (2009).

45. F. Corrêa, J. Key, B. Kuhlman, Kevin H. Gardner, Computational Repacking of HIF-2α Cavity Replaces Water-Based Stabilized Core. Structure 24, 1918–1927 (2016).

46. X. Xu et al., Identification of small-molecule ligand-binding sites on and in the ARNT PAS-B domain. J Biol Chem 300, 107606 (2024).

47. J. Rutter, C. H. Michnoff, S. M. Harper, K. H. Gardner, S. L. McKnight, PAS kinase: An evolutionarily conserved PAS domain-regulated serine/threonine kinase. Proceedings of the National Academy of Sciences 98, 8991–8996 (2001).

48. A. Möglich, R. A. Ayers, K. Moffat, Structure and Signaling Mechanism of Per-ARNT-Sim Domains. Structure 17, 1282–1294 (2009).

49. V. Gonzalez et al., The partitioned Rhizobium etli genome: Genetic and metabolic redundancy in seven interacting replicons. Proceedings of the National Academy of Sciences 103, 3834–3839 (2006).

50. F. Correa, W. H. Ko, V. Ocasio, R. A. Bogomolni, K. H. Gardner, Blue light regulated two-component systems: enzymatic and functional analyses of light-oxygen-voltage (LOV)-histidine kinases and downstream response regulators. Biochemistry 52, 4656–4666 (2013).

51. G. Rivera-Cancel, W. H. Ko, D. R. Tomchick, F. Correa, K. H. Gardner, Full-length structure of a monomeric histidine kinase reveals basis for sensory regulation. Proc Natl Acad Sci U S A 111, 17839–17844 (2014).

52. A. Vulpetti, U. Hommel, G. Landrum, R. Lewis, C. Dalvit, Design and NMR-Based Screening of LEF, a Library of Chemical Fragments with Different Local Environment of Fluorine. J Am Chem Soc 131, 12949–12959 (2009).

53. T. M. Jacobs et al., Design of structurally distinct proteins using strategies inspired by evolution. Science 352, 687–690 (2016).

54. J. Feng et al., A general strategy to construct small molecule biosensors in eukaryotes. Elife 4, e10606 (2015).

55. D. Sitkoff, D. A. Case, Theories of chemical shift anisotropies in proteins and nucleic acids. Progress in Nuclear Magnetic Resonance Spectroscopy 32, 165–190 (1998).

56. M. Beck Erlach et al., Pressure dependence of side chain 13C chemical shifts in model peptides Ac-Gly-Gly-Xxx-Ala-NH2. J Biomol NMR 69, 53–67 (2017).

57. R. Kitahara, K. Hata, H. Li, M. P. Williamson, K. Akasaka, Pressure-induced chemical shifts as probes for conformational fluctuations in proteins. Progress in Nuclear Magnetic Resonance Spectroscopy 71, 35–58 (2013).

58. R. J. Quinlan, G. D. Reinhart, Baroresistant buffer mixtures for biochemical analyses. Anal Biochem 341, 69–76 (2005).

59. B. A. Johnson, From Raw Data to Protein Backbone Chemical Shifts Using NMRFx Processing and NMRViewJ Analysis. Methods Mol Biol 1688, 257–310 (2018).

60. B. A. Johnson, R. A. Blevins, NMRView: a computer program for the visualization and analysis of NMR data. J Biomol NMR 4, 603–614 (1994).

61. E. Koag et al., NMRFx: Integrated Software for NMR Data Processing, Visualization, Analysis and Structure Calculation. bioRxiv 10.1101/2025.08.26.672401 (2025).

62. M. Norris, B. Fetler, J. Marchant, B. A. Johnson, NMRFx Processor: a cross-platform NMR data processing program. J Biomol NMR 65, 205–216 (2016).

