## Supplementary Information for "Use of High Pressure NMR Spectroscopy to Rapidly Identify Proteins with Internal Ligand-Binding Voids"

**This PDF file includes:**

SI Methods

Tables S1, S2

Figs. S1 to S10

SI References

**SI Materials and Methods**

*Protein preparation* Human HIF-2α PAS-B (237-350) (1), *Calypte anna* HIF-2α caEPAS1 PAS-B (231-341), *Danio rerio* HIF- 2α drEPAS1 PAS-B (237-346), human ARNT PAS-B (356-470) (2), human PASK PAS-A (131-285) (3), GB1 (4) and human HIF-2α PAS-B Design1 (5) were expressed in *E. coli* BL21(DE3) using ^15^NH_4_Cl-containing M9 minimal media and purified as previously described.

RE137 is an N-terminal 137 aa fragment of the *Rhizobium etli* histidine kinase RE356 (Uniprot accession ID: Q2KD32, from genome reported in refs. (6-8)), including a single PAS domain. Genomic DNA encoding this fragment was cloned into the pHis-Gβ1-parallel expression vector (9), which was subsequently transformed into *E. coli* BL21(DE3) cells which were subsequently grown at 37 °C in M9 minimal media supplemented with 1 g/L ^15^NH_4_Cl and 100 μg/mL of ampicillin. Cells were grown to an OD_600_ of 0.6–0.8 prior to induction with 100 µM isopropylthiogalactoside (IPTG) at 18°C and harvested 18 hr post induction. The harvested pellet was suspended in 50 mM Tris (pH 7.5), 100 mM NaCl buffer and sonicated at 4°C. The supernatant was separated by centrifugation and subjected to Ni-Sepharose affinity purification, with the pHis-Gβ1-RE137 protein obtained by gradient elution with 5–500 mM imidazole in the same buffer. Eluted samples were subsequently digested with His_6_-TEV protease (10) overnight at 4°C to separate the affinity tag from the RE137 protein. The cleaved protein was subjected to a second round of Ni^2+^ affinity chromatography, this time collecting the RE137 protein found in the flow-through fraction. These fractions were concentrated (Amicon Ultra, Millipore) and subjected to a final purification step via Superdex 75 size exclusion chromatography in 50 mM Tris (pH 7.5), 100 mM NaCl, 5 mM DTT buffer. Fractions corresponding to the dimeric RE137 were concentrated, flash frozen in liquid N_2_ and stored at -80 °C.

For the designed protein CA01 (1-114), an overexpression plasmid construct (1) was transformed into *E. coli* BL21(DE3) and subsequently grown at 37 °C in M9 minimal media supplemented with 1 g/L ^15^NH_4_Cl and 50 μg/mL of ampicillin. Cells were grown until the OD_600_ reached approximately 0.8 prior to induction with 660 μM of IPTG at 25°C and harvested after 15 hours of protein expression. The harvested pellet was resuspended in 20 mM Tris-HCl (pH 8.0), 300 mM NaCl, 80 mM imidazole buffer and sonicated at 4 °C. The supernatant was separated by centrifugation and subjected to Ni-Sepharose affinity purification to elute the pHis-SUMO-CA01 protein with 500 mM imidazole in the same buffer. The purified protein was then dialyzed overnight into 4L of the same buffer without imidazole, before being cleaved with SUMO protease (2) overnight at 4 °C or 1 hour at room temperature. Post-cleavage, samples were again applied to the Ni-Sepharose affinity column equilibrated with dialysis buffer and the flow-through was collected as the final sample and buffer exchanged into a baroresistant buffer of 35 mM Tris (pH 7.5), 14.7 mM phosphate, 50 mM NaCl, 5 mM DTT (3). The purity of the protein was assessed via SDS-PAGE electrophoresis.

The human bromodomain (Brd) proteins Brd2-D1 (73-194) (11), Brd2-D2 (344-455) (12), Brd3-D2 (306-415) (13), Brd4-D1 (42-168) (14), Brd4-D2 (333-460) (12), and PBRM1 (43-154) (15) plasmid constructs were transformed into *E. coli* BL21(DE3) and subsequently grown at 37°C in M9 minimal media supplemented with 1 g/L ^15^NH_4_Cl and 100 μg/mL of ampicillin. Cells were grown to an OD_600_ of ~0.8 prior to induction with 1 mM IPTG at 17°C and harvested 18 hours post induction. The harvested pellet was suspended in 50 mM Tris (pH 7.5), 100 mM NaCl, 1 mM DTT buffer and sonicated at 4°C. The supernatant was separated by centrifugation and subjected to Ni-Sepharose affinity purification to elute the pHis-TEV-Brd protein with 500 mM imidazole in the same buffer. Eluted samples were subsequently digested with His_6_-TEV protease (9) overnight at 4 °C to separate the affinity tag from the protein. The cleaved protein was subjected to a second round of Ni-Sepharose chromatography, this time collecting the Brd protein found in the flow-through fraction. These fractions were concentrated (Amicon Ultra, Millipore) and subjected to final step of purification via Superdex 75 size exclusion chromatography in 50 mM Tris (pH 7.5), 50 mM NaCl, 1 mM DTT buffer. Fractions corresponding to the Brd protein were concentrated, flash frozen in liquid N_2_ and stored at -80 °C.

His-tagged versions of the rat fatty acid-binding proteins (FABPs), liver type L-FABP (1-127) and intestinal type I-FABP (1-133), were purified as previously described (16). Plasmid constructs were transformed into BL21(AI) cells and subsequently grown at 37 °C overnight in M9 minimal media supplemented with 1 g/L ^15^NH_4_Cl. The cells were grown until the OD_600_ reached approximately 0.8 prior to induction with 2.0% (g/v) L-arabinose (Sigma-Aldrich, St. Louis, MO) and harvested after 4 hours of protein expression. The harvested pellet was re-suspended in 20 mM Tris-HCl (pH 8), 300 mM NaCl, 1 mM PMSF, 0.5 mg of DNase, 50 µM of MgSO_4_, pH 8. The supernatant was separated by centrifugation and subsequently loaded onto a 5 mL HiTrap HP Ni column before being washed extensively with buffer (20 mM Tris-HCl pH 8, 300 mM NaCl, 25 mM imidazole). Bound protein was then treated with TEV protease (1.5 mg TEV added on column and incubated 24-48 hr at room temperature) to remove the His-tag; cleaved protein eluted in the unbound fractions. Copurifying lipids were removed for L-FABP by extraction with n-butanol, using three extractions against 1/3 volumes of n-butanol (15’ apiece, retaining aqueous protein-containing phase); for I-FABP, lipids were removed by 3 hr incubation with pre-swelled HAP-Dextran at 37 °C and eluted with buffer (16). Post-extraction, protein samples were concentrated and buffer-exchanged by centrifugal filtration; removal of lipids was confirmed by comparing ^1^H-^15^N HSQC spectra to previously published spectra reference spectra (17).

For all proteins, concentration was estimated using the theoretical extinction coefficient based on amino acid sequence and calculated by ProtParam (ExPASy) (18). All samples used for subsequent analyses were > 95% pure as determined by SDS-PAGE and MALDI-TOF mass spectrometry.

*AlphaFold3 and Boltz-2 Model Generation* To generate structural models of the RE137 PAS domain, we input residues 1-137 of the *Rhizobium etli* histidine kinase RE356 sequence (8) (UniProt accession ID: Q2KD32) into AlphaFold3 (AF3) (19-21) high-accuracy prediction on Google Colab Jupyter Notebook. The model shown in **Fig. 5** was ranked best among five generated models, as assessed with the pLDDT confidence measure. This model has been deposited for the apo RE137 is available in ModelArchive ([modelarchive.org)](https://nam02.safelinks.protection.outlook.com/?url=https%3A%2F%2Furldefense.com%2Fv3%2F__https%3A%2F%2Fmodelarchive.org%2Fdoi%2F10.5452%2Fma-bi5qr__%3B!!GekbXoL5ynDpFgM!Xq3oXvpA22t3aWCmlTKpzf3r1nceOzihRs0Ck7R6bMOmnOkYyloJYZiaw_dUmuFjNNh3JLYrSjPKDdjg9FiJNysBN45GyOg%24&data=05%7C01%7Crazad%40gc.cuny.edu%7C61c78435b376492640d108dbbed88bed%7C6f60f0b35f064e099715989dba8cc7d8%7C0%7C0%7C638313609561315354%7CUnknown%7CTWFpbGZsb3d8eyJWIjoiMC4wLjAwMDAiLCJQIjoiV2luMzIiLCJBTiI6Ik1haWwiLCJXVCI6Mn0%3D%7C3000%7C%7C%7C&sdata=kPr35RwPTr0ThuPXqyMdu8wRKgc9R5KEIkEpzMMMhk8%3D&reserved=0) (22) (ID: ma-ev4h3).

To model the RE137 PAS domain in complex with compounds 15 and 47, and hPASK PAS-A with compounds KG-535 and KG-571, we used the Boltz-2 (23) structure prediction model for protein ligand complexes. Protein sequences and ligand SMILES strings were input into the publicly available Boltz2 inference pipeline on Google Colab and GitHub, using default parameters. The resulting models were selected from the five outputs based on pocket-bound poses with minimal steric clash and plausible geometry. Ligand-binding residues were defined as those with atoms within 4 Å of the ligand (**Figs. 5**, **S10**). These models have been deposited and are available in ModelArchive ([modelarchive.org)](https://nam02.safelinks.protection.outlook.com/?url=https%3A%2F%2Furldefense.com%2Fv3%2F__https%3A%2F%2Fmodelarchive.org%2Fdoi%2F10.5452%2Fma-bi5qr__%3B!!GekbXoL5ynDpFgM!Xq3oXvpA22t3aWCmlTKpzf3r1nceOzihRs0Ck7R6bMOmnOkYyloJYZiaw_dUmuFjNNh3JLYrSjPKDdjg9FiJNysBN45GyOg%24&data=05%7C01%7Crazad%40gc.cuny.edu%7C61c78435b376492640d108dbbed88bed%7C6f60f0b35f064e099715989dba8cc7d8%7C0%7C0%7C638313609561315354%7CUnknown%7CTWFpbGZsb3d8eyJWIjoiMC4wLjAwMDAiLCJQIjoiV2luMzIiLCJBTiI6Ik1haWwiLCJXVCI6Mn0%3D%7C3000%7C%7C%7C&sdata=kPr35RwPTr0ThuPXqyMdu8wRKgc9R5KEIkEpzMMMhk8%3D&reserved=0) (22). Structural models of the RE137 PAS domain in complex with compound 15 (ID: ma-irsym) and compound 47 (ID: ma-egwby), and the hPASK PAS-A domain in complex with compound KG-535 (ID: ma-blpib) and compound KG-571 (ID: ma-awqgk).

### NMR data acquisition All NMR experiments were carried out on Bruker Avance III HD NMR spectrometers at 700 and 800 MHz equipped with 5 mm cryoprobes (QCI-F with pulsed-field Z gradient (700 MHz) or TCI with pulsed-field XYZ gradients (800 MHz)), and Topspin 3.5 software (Karlsruhe, Germany). Pressure NMR experiments were conducted with 100-700 µM protein in a baroresistant buffer containing a pair of buffer compounds (14.7 mM Tris pH 7.4, 35 mM sodium phosphate pH 7.4, 20 mM NaCl, and 20% D_2_O) to limit pressure-induced pH changes (24). For protein/ligand complexes, the selected small molecule was added (to final concentrations indicated in each figure legend) and incubated at 298 K for at least 30 min before data acquisition.

Samples were transferred to a zirconia tube of 3/5 mm inner/outer diameter connected to a syringe pump (Xtreme 60 pump, Daedalus Innovations LLC, Aston, PA). A layer of mineral oil (270 μL) was added to separate the sample from the hydrostatic pressure. Sensitivity-enhanced ^1^H/^15^N HSQC experiments were acquired at 298 K using spectral widths of 40 and 16 ppm in the t_1_ (^15^N) and t_2_ (^1^H) dimensions, respectively. As shown in **Fig. 2**, spectra were acquired with pressure steps between 20-2500 bar, interleaving experiments at high pressure and 20 bar to establish reversibility and allowing 10 min between pressure changes before acquiring data. A complete list of all proteins and complexes analyzed in this way is provided on **Table S1**.

*NMR data analysis – high pressure titration* All NMR data were processed and analyzed with NMRViewJ (25, 26) with NMRFx (25, 27, 28). After individually processing each 2D ^1^H/^15^N HSQC spectrum, we picked peaks and tracked their changes in chemical shift as a function of pressure. Separately handling movements in the ^1^H and ^15^N dimensions, we fit these trends to the following second-order polynomial equation (29):

$\delta_{i}=a_{i}+b_{i}p+c_{i}p^{2}$ **(Eq. S1)**

where p is the pressure (bar), δ*_i_* is the chemical shift for the *i*th residue, a*_i_* is the chemical shift at 20 bar, and *b_i_* and c*_i_* are the coefficients for the linear and non-linear effects of pressure on chemical shift.

Residue-specific b*_i_* and c*_i_* values for seven reference proteins in **Fig. 3** (BPTI, GB1, lysozyme, RalFree, HPr, BlgB, and RalComplex) were generously provided by Prof. Kazuyuki Akasaka (29, 30).

*Volume calculations* Two types of volumes were calculated for various proteins: the volume of specific, single cavities within a protein (“cavity volume”) and the volume summing all voids distributed throughout a structure (“total void volume”).

For cavity volume calculations, we utilized *cavfinder* (31), an in-house Python script which uses a grid-based search approach to identify internal cavities and quantitate their volumes. Reported cavity volumes correspond to the volume of the largest single cavity within a protein.

For total void volume calculations, we used *ProteinVolume* (32), which calculates the sum of all voids, cavities and packing defects within a protein by calculating the difference in volume between the solvent-excluded geometric volume (calculated with a Lee-Richards rolling sphere approach) and the van der Waals volume (calculated from the volume of all amino acids in the sequence). While this approach does not identify spatial locations of such voids, the “total void volume” represents their collective contribution throughout the protein as a whole. For each calculation, the energy minimization function was activated. A starting (ending) probe size of 0.08 (0.02) Å^3^, with a surface minimum distance of 0.1 Å^3^ were selected. The list of void volume determination for each protein in Fig. 3 are (PDB entries): HIF-2α PAS-B (3F1P, chain A), ARNT PAS-B (4EQ1, chain B), hPASK PAS-A (1LL8), GB1 (2GB1), BPTI (5PTI), blgb (1CJ5), lysozyme (1E8L), HPr (1QR5), RalFree (1LXD), RalComplex (1LFD).

*NMR based fragment screening – RE137* To identify small molecule binders of the RE137 PAS domain, we screened a small in-house library composed of 58 fluorinated members of the Prestwick library. These compounds were assembled into 5 mixtures of 11-12 compounds for an initial screen, grouping compounds with non-overlapping ^19^F NMR signals to facilitate ligand-based screening approaches. For a screen, each mixture (containing 100 µM of each ligand; a total of 1.2% DMSO-d_6_ working concentration) had 20 µM RE137 added, followed by 1D ^19^F R_2_ filter screens. Hits were verified by assembling 1:1 mixture of individual compounds with U-^15^N RE137 (40 µM each) and acquiring both ligand- and protein-detected spectra (1D ^19^F R_2_ filter, SF ^1^H/^15^N HSQC).

*Crystallization, Structure Determination, and Refinement – ARNT PAS-B co-crystallized with KG-548* Purified ARNT PAS-B WT protein was exchanged into a buffer of 25 mM Tris (pH 7.5), 17 mM NaCl, and 5 mM beta-mercaptoethanol (BME). Sample was concentrated to 10 mg/ml (~ 720 µM) and mixed with KG-548 stock to reach a final ligand concentration of 5 mM (maximum soluble concentration) and 2% DMSO. The sample was allowed to equilibrate for 1 hr on ice and set up for co-crystallization at RT using the sitting drop vapor diffusion method. Protein crystallization screening was set up using the following sparse matrix screens (Qiagen) – JCSG+, PEGs, Classics, PACT and the Protein Complex suites. The best diffracting crystals were obtained with the Protein Complex suite, condition H6 at a ratio of 2:1 (1.6 M magnesium sulfate, 0.1 M MES, pH 6.5; 0.4 µL protein and 0.2 µL of precipitant solution mix). Diffraction data were obtained on 24-ID-E at the Advanced Light Source (Argonne National Laboratory, Argonne, IL), and AMX 17-ID1 at National Synchrotron Light Source II (Brookhaven National Laboratory, Upton, NY). Data were processed with autoPROC (33). The structure of ARNT PAS-B/KG-548 was solved via molecular replacement with apo ARNT PAS-B (PDB: 4EQ1) as a search model using Phaser (29, 30). The structure was iteratively evaluated and manually corrected in Coot and refined with phenix.refine (34, 35). The model was optimized with PDB-REDO before the deposition (36). Data collection and refinement information are presented in **Table S2**.

| **PAS DOMAINS** | | | **FABPs / BROMODOMAINS / ENGINEERED** | | | **AKASAKA STANDARDS** | | |
| --- | --- | --- | --- | --- | --- | --- | --- | --- |
| **protein** | **^1^H** | **^15^N** | **protein** | **^1^H** | **^15^N** | **protein** | **^1^H** | **^15^N** |
| HIF-2α PAS-B (37) | 72 | 72 | L-FABP (16) | 95 | 95 | GB1’ | 61 | 61 |
| HIF-2α PAS-B / 2 (31) | 85 | 85 | L-FABP / oleate (16) | 100 | 100 | GB1  (29, 30) | 55 | 55 |
| HIF-2α PAS-B / 37 (38) | 88 | 88 | I-FABP (16) | 95 | 95 | BPTI  (29, 30) | 52 | 52 |
| HIF-2α D1 (5) | 71 | 71 | I-FABP / oleate (16) | 108 | 108 | BlgB  (29, 30) | 126 | 125 |
| HIF-2α PAS-B M289I | 56 | 56 |  |  |  | HPr  (29, 30) | 84 | 85 |
| HIF-2α PAS-B S304T / 2 | 75 | 75 | Brd2-D1 (11) | 125 | 125 | Ral Free (29, 30) | 79 | 79 |
| HIF-2α PAS-B S304M (39) | 83 | 83 | Brd2-D2 (12) | 98 | 98 | Ral (29, 30) | 71 | 71 |
| caEPAS1 PAS-B | 75 | 75 | Brd2-D2 / MS417 (12) | 52 | 52 | Lysozyme  (29, 30) | 126 | 126 |
| caEPAS1 PAS-B / 2 | 103 | 103 | Brd3-D2 (13) | 104 | 104 |  |  |  |
| drEPAS1 PAS-B | 81 | 81 | Brd4-D1 (14) | 109 | 109 |  |  |  |
| ARNT PAS-B (2) | 92 | 92 | Brd4-D2 (12) | 90 | 90 |  |  |  |
| ARNT PAS-B / KG-548 (40) | 82 | 82 | PBRM1 (15) | 109 | 109 |  |  |  |
| ARNT PAS-B / KG-655 (40) | 84 | 84 |  |  |  |  |  |  |
| ARNT PAS-B Y456T (41) | 49 | 49 | CA01 (42) | 100 | 100 |  |  |  |
| ARNT PAS-B F444Q/F446A/Y456T (41) | 90 | 90 |  |  |  |  |  |  |
| hPASK PAS-A (3) | 76 | 76 |  |  |  |  |  |  |
| hPASK PAS-A / KG-535 (3) | 77 | 77 |  |  |  |  |  |  |
| hPASK PAS-A / KG-571 (3) | 74 | 74 |  |  |  |  |  |  |
| RE137 (8) | 30 | 30 |  |  |  |  |  |  |
| RE137 / 15 (8) | 24 | 24 |  |  |  |  |  |  |
| RE137 / 47 (8) | 21 | 21 |  |  |  |  |  |  |
| EL222 / FMN (43) | 116 | 116 |  |  |  |  |  |  |

**Table S1: A total of 42 proteins / protein-ligand complexes were used for these analyses. Columns from left to right indicate:**

**PAS domains**, chiefly include PAS-B domains from the human HIF-2 (HIF-2α/ARNT) complex, with several variants bound to small molecule inhibitors or containing point mutations. Two HIF-2α PAS-B homologs from non-human vertebrates were also tested, caEPAS1 (*Calypte anna* EPAS1 [= hummingbird HIF-2α]), and drEPAS1 (*Danio rerio* EPAS1 [= zebrafish HIF-2α]). Other PAS domains were obtained from human PAS kinase (hPASK), the bacterial *R. etli* histidine kinase described above, and the *E. litoralis* EL222 transcription factor.

**FABPs/Bromodomains/Engineered**, includes a mix of liver and intestinal fatty acid binding proteins (L- and I-FABP, respectively) on their own and bound to oleate; six bromodomains, including one tested as an apo- protein and bound to the MS417 inhibitor; and the Rosetta-designed CA01 artificial helical bundle.

**Akasaka standards**, indicating datasets provided by K. Akasaka from analyses previously reported in cited publications. Note that the first entry (GB1’) was independently expressed and analyzed in the Gardner laboratory for comparisons with the previously acquired data in the Akasaka lab.

Samples of **protein/small molecule complexes** all contained a minimum of 250 µM small molecule ligand.

| **Beamline** | NSLS-II BEAMLINE 19-ID | **Solvent** | 3 |
| --- | --- | --- | --- |
| **Wavelength** | 0.979 | **Protein residues** | 212 |
| **Resolution range** | 69.4 - 1.97  (2.04 - 1.97) | **RMS (bonds)** | 0.011 |
| **Space group** | P 31 2 1 | **RMS (angles)** | 1.30 |
| **Unit cell** | 80.135 80.135 100.718 90 90 120 | **Ramachandran favored (%)** | 99.03 |
| **Unique reflections** | 27006 (2701) | **Ramachandran allowed (%)** | 0.97 |
| **Completeness (%)** | 99.76 (98.15) | **Ramachandran outliers (%)** | 0.00 |
| **Wilson B-factor** | 61.64 | **Rotamer outliers (%)** | 2.99 |
| **Reflections used in refinement** | 27006 (2651) | **Clash score** | 2.57 |
| **Reflections used for R-free** | 1977 (198) | **Average B-factor (all atoms)** | 79.07 |
| **R-work** | 0.2064 (0.4268) | **Average B-factor (macromolecules)** | 79.23 |
| **R-free** | 0.2344 (0.4054) | **Average B-factor (ligand)** | 67.79 |
| **Number of non-hydrogen atoms** | 1787 | **Average B-factor (solvent)** | 57.66 |
| **Number of macromolecule atoms** | 1765 | **Number of TLS groups** | 2 |
| **Number of ligand atoms** | 19 | **PDB accession code** | 8G4A |

*Statistics for the highest-resolution shell are shown in parentheses.*

**Table S2: Data collection and refinement statistics for ARNT PAS-B:KG-548 complex X-ray crystal structure.**

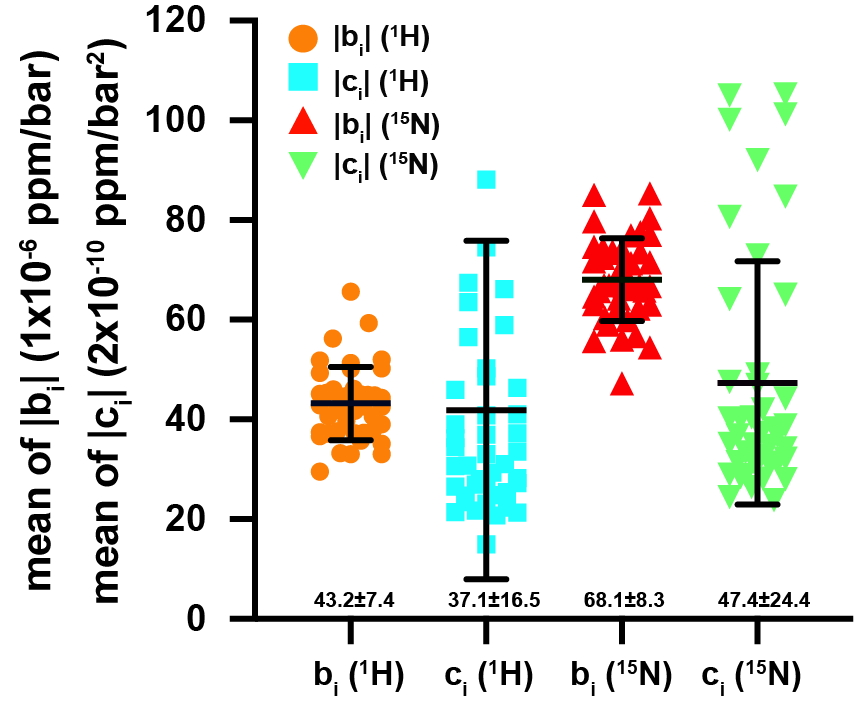

**Fig. S1:** **Distribution of the linear and non-linear coefficients of pressure-dependent chemical shift changes for 42 proteins and protein/ligand complexes (listed in Table S1), summarizing information from over 3400 amide trajectories.** Each point represents the mean of the absolute value for each listed parameter, including data for all residues within a given protein. Parameters represented are the linear |b_i_| component for the amide ^1^H (orange) and ^15^N (red) nuclei, along with the nonlinear |c_i_| components for the amide ^1^H (cyan) and ^15^N (green) nuclei. The mean and standard deviation of these values are identified by black lines, and are reported underneath as mean ± s.d.

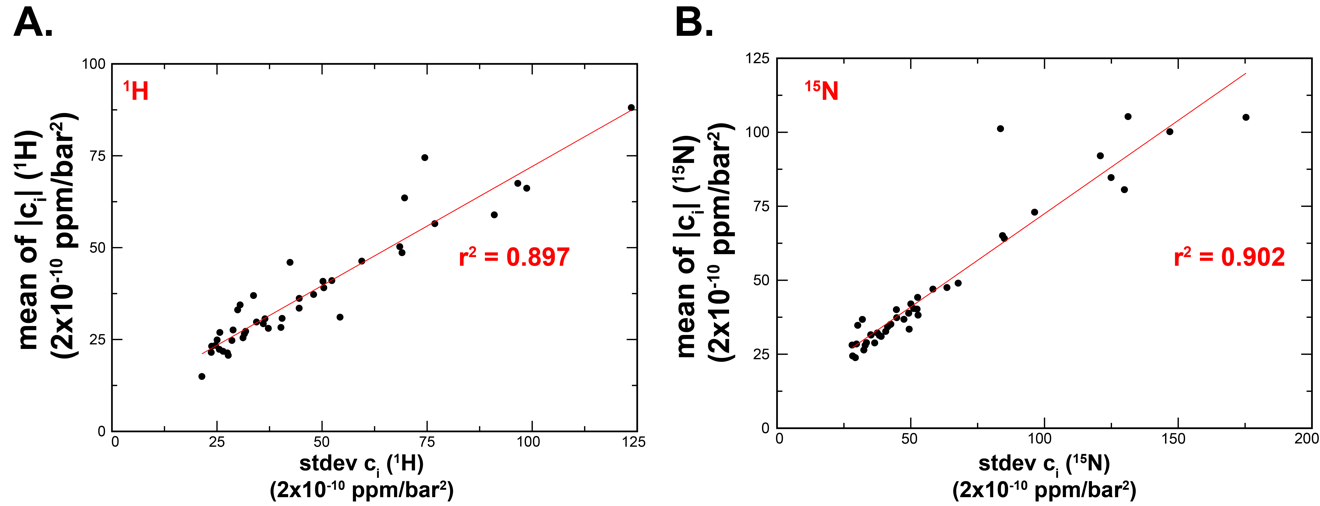

**Fig. S2:** **Correlation between different representations of the diversity of nonlinear pressure-dependent chemical shift coefficients (c*_i_*) for backbone amides.** Graphs show correlations between the standard deviation of c*_i_* values (stdev(c*_i_*), x-axes) and means of the absolute values of c*_i_* values (y-axes, as previously established by ref. (30)), using separate analyses for ^1^H (panel A) and ^15^N (panel B). Each point summarizes the individual residue-specific values for the proteins listed in Fig. S1. Both ^1^H and ^15^N analyses show excellent correlations between the two types of analyses, with correlation coefficients for linear regression fits (r^2^) of 0.897 and 0.902 for the amide proton and nitrogen, respectively.

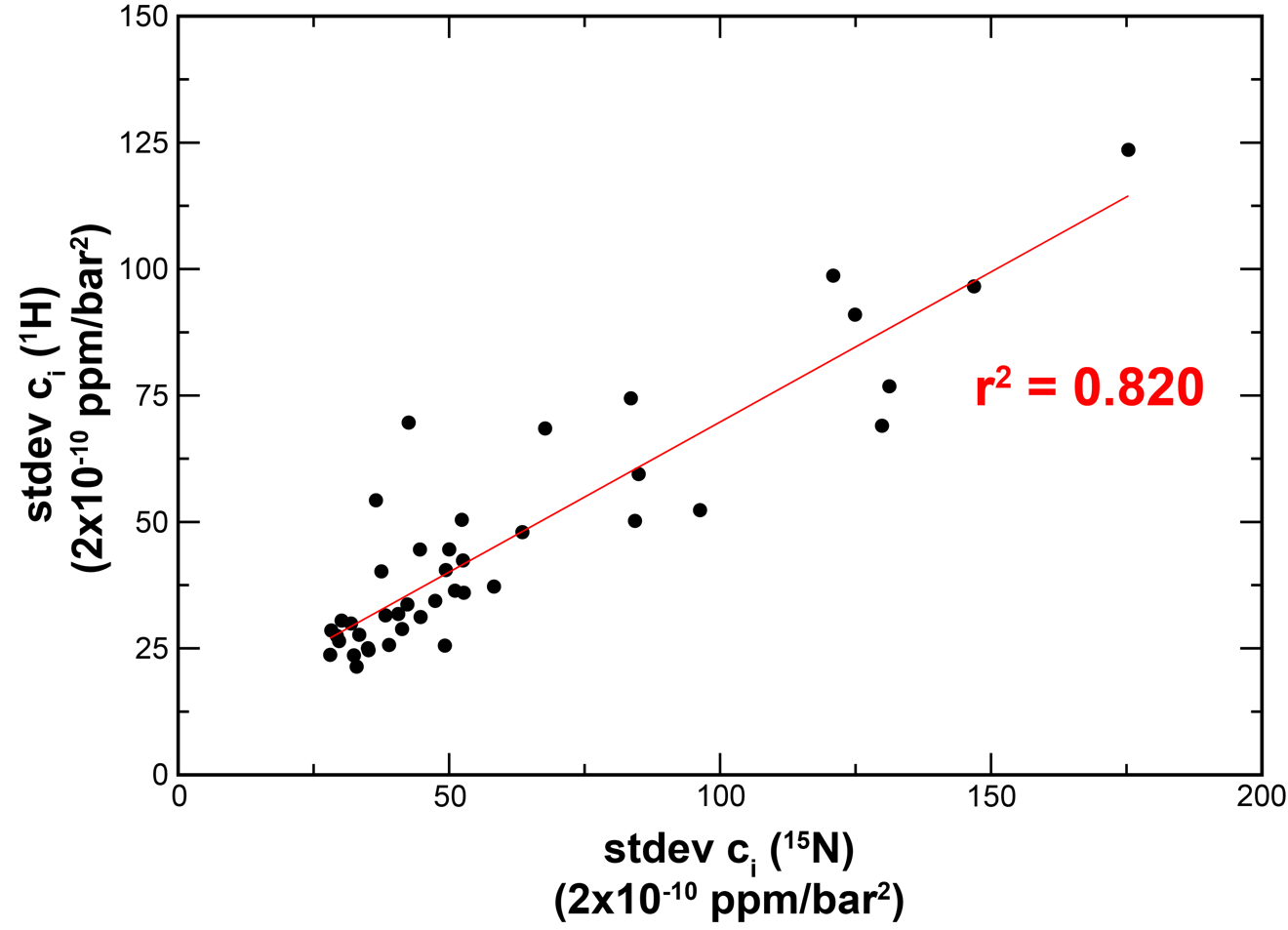

**Fig. S3: Correlation between the diversity of nonlinear pressure-dependent chemical shift coefficients (c*_i_*) for backbone amide ^1^H and ^15^N sites.** Plot showing the stdev(c*_i_*) values for ^1^H versus ^15^N, with a correlation coefficient for the linear regression fit (r^2^) of 0.820. Each point summarizes the individual residue-specific values for the proteins listed in **Fig. S1.**

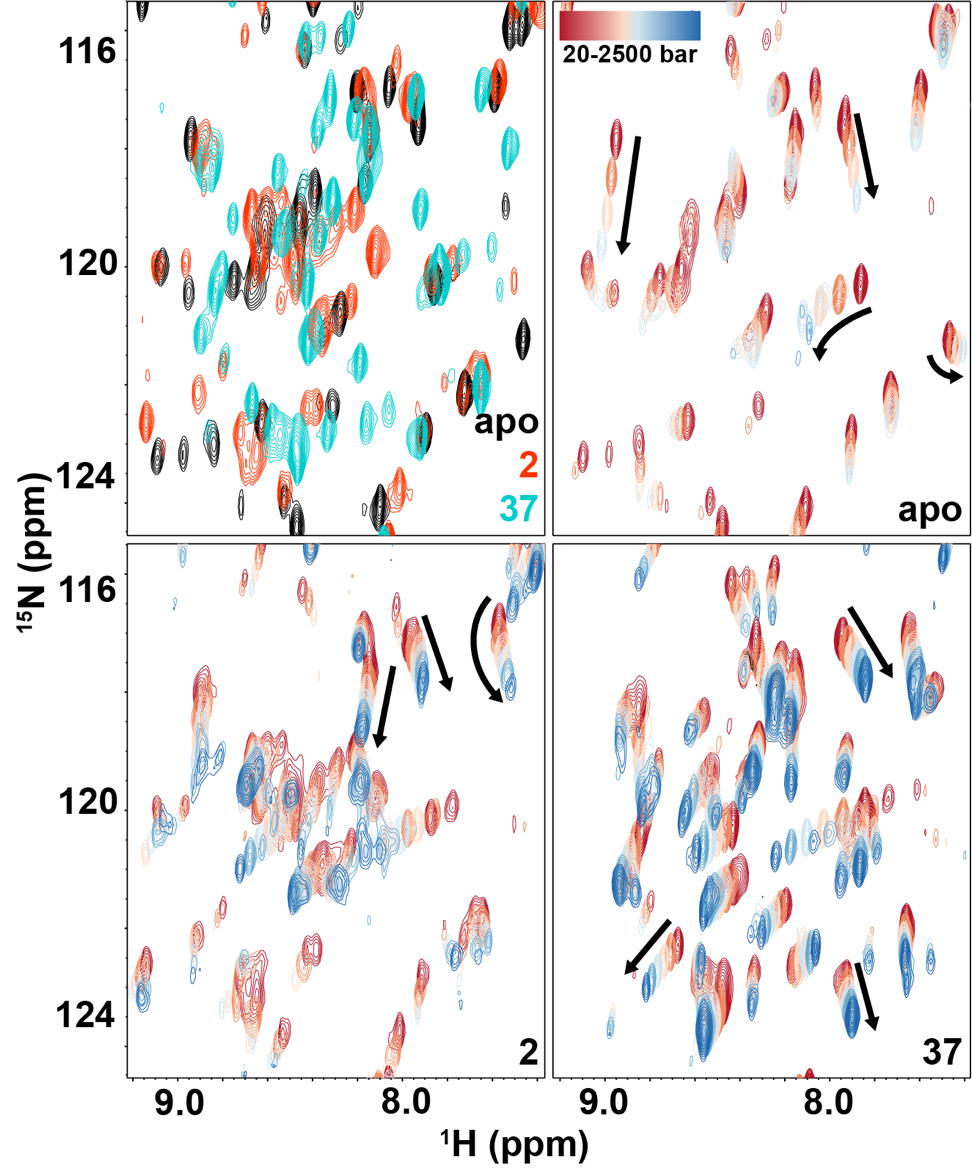

**Fig. S4: Spectra showing the ligand binding and pressure dependence of HIF-2**α **PAS-B in apo- and ligand-bound forms.** (top left) Overlay of ^1^H/^15^N-HSQC spectra of HIF-2α PAS-B in the apo- (black), 2-bound (red) and 37-bound (cyan) forms at 1 atm (1.01 bar). (other panels) Overlays of ^1^H/^15^N-HSQC spectra acquired under increasing pressure from 20-2500 bar (red-to-blue) of HIF-2α PAS-B apo- (top right), 2-bound (bottom left) and 37-bound (bottom right) forms. Selected trajectories are indicated by arrows to show direction of pressure-induced chemical shift changes. All samples contained 280 µM HIF-2α PAS-B protein, with 310 µM compounds added when indicated.

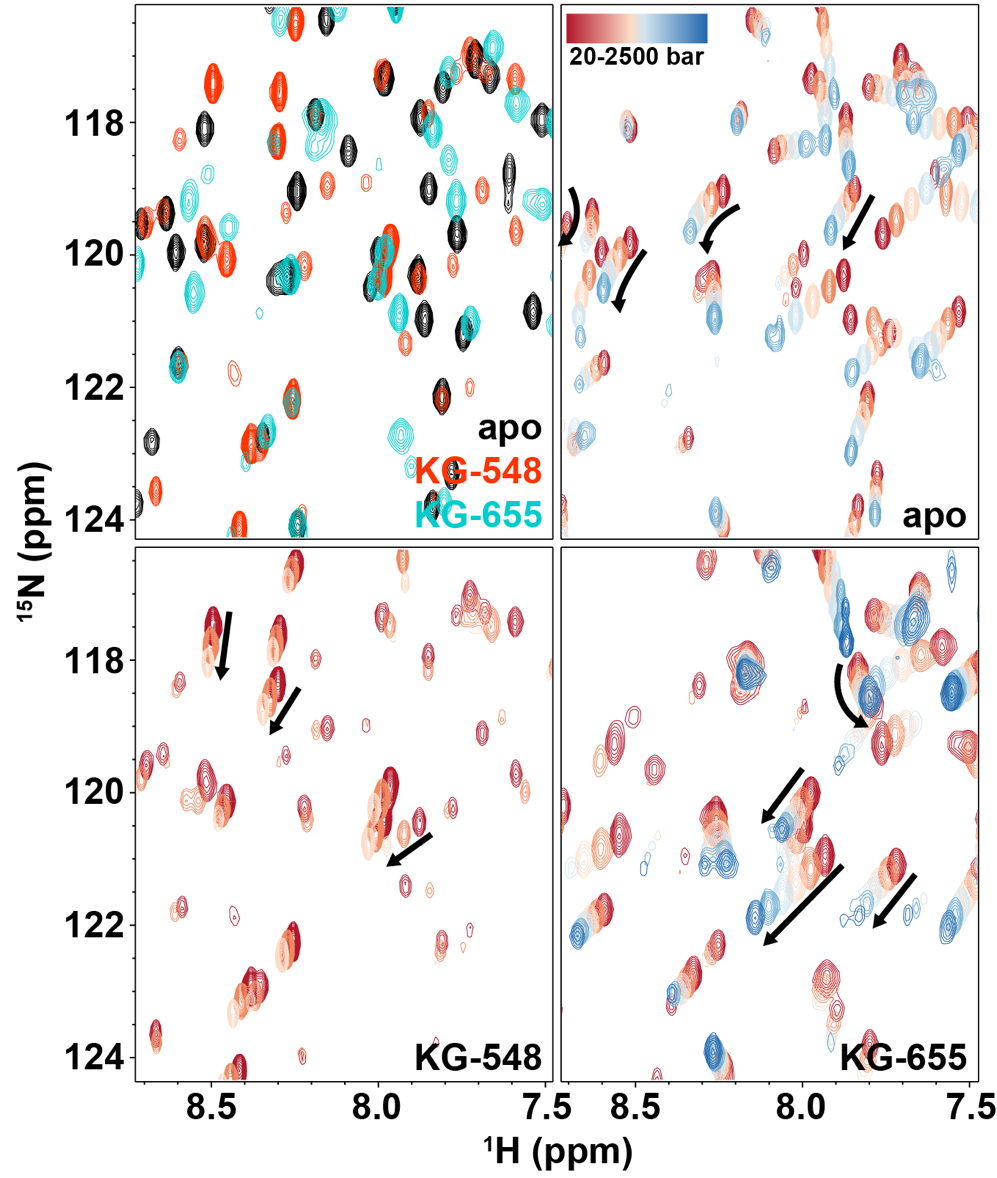

**Fig. S5: Spectra showing the ligand binding and pressure dependence of ARNT PAS-B in apo- and ligand-bound forms.** (top left) Overlay of ^1^H/^15^N-HSQC spectra of ARNT PAS-B in its apo- (black), KG-548-bound (red) and KG-655-bound (cyan) forms. (other panels) Overlays of ^1^H/^15^N-HSQC spectra acquired under increasing pressure from 20-2500 bar (red-to-blue) of ARNT PAS-B apo- (top right), KG-548-bound (bottom left) and KG-655-bound (bottom right) forms. Selected trajectories are indicated by arrows to show direction of pressure-induced chemical shift changes. Samples contained either 250 µM ARNT PAS-B protein, 300 µM ARNT PAS-B with 300 µM KG-548, or 250 µM ARNT PAS-B with 250 µM KG-655.

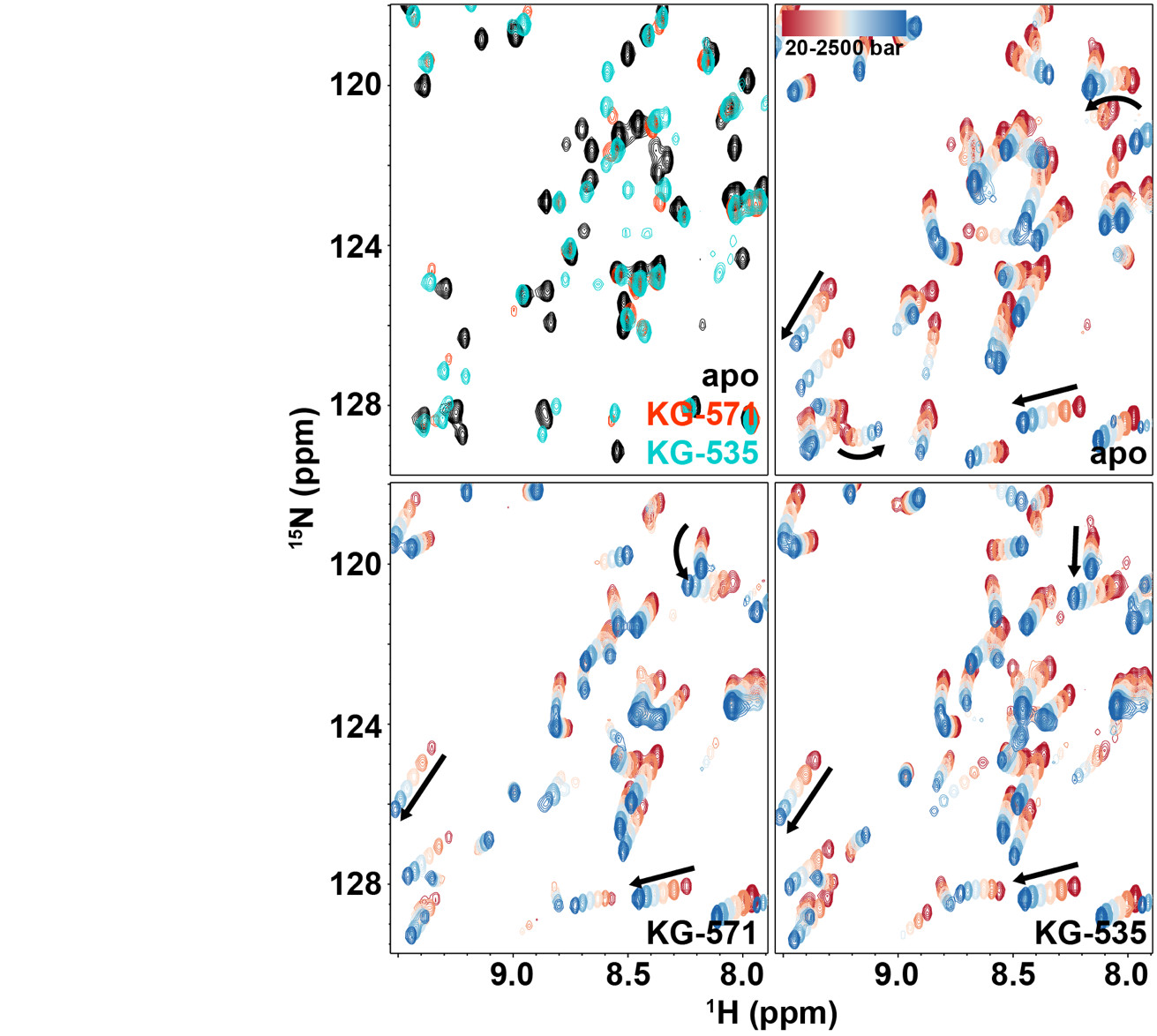

**Fig. S6:** **Spectra showing the ligand binding and pressure dependence of hPASK PAS-A in apo- and ligand-bound forms.** (top left) Overlay of ^1^H/^15^N-HSQC spectra of hPASK PAS-A in its apo- (black), KG-571-bound (red) and KG-535-bound (cyan) forms. (other panels) Overlays of ^1^H/^15^N-HSQC spectra acquired under increasing pressure from 20-2500 bar (red-to-blue) of hPASK PAS-A apo (top right), KG-571-bound (bottom left) and KG-535-bound (bottom right) forms. Selected trajectories are indicated by arrows to show direction of pressure-induced chemical shift changes. All samples contained 675 µM hPASK PAS-A protein, with 1 mM compounds added when indicated.

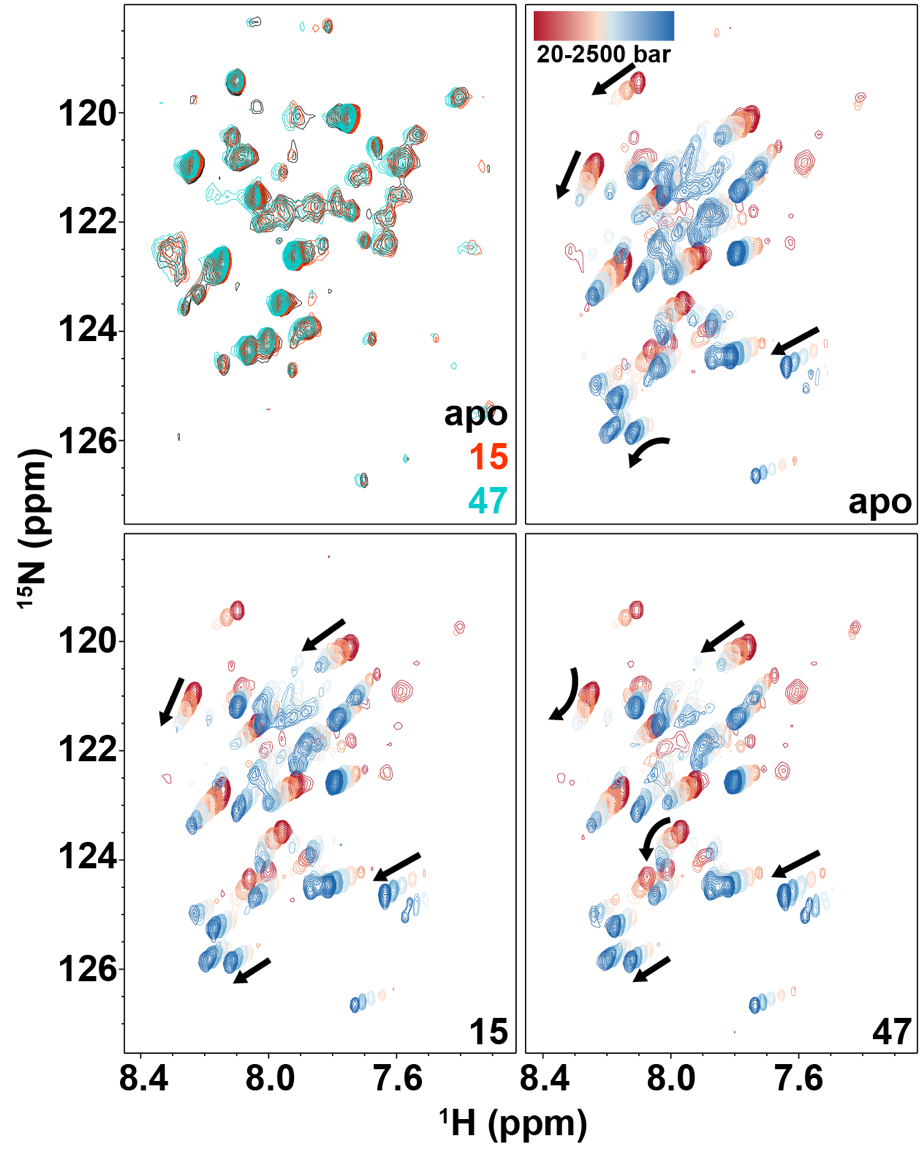

**Fig. S7: Spectra showing the ligand binding and pressure dependence of RE137 in apo- and ligand-bound forms.** (top left) Overlay of ^1^H/^15^N-HSQC spectra of RE137 in its apo (black), 15-bound (red) and 47-bound (cyan) forms. (other panels) Overlays of ^1^H/^15^N-HSQC spectra acquired under increasing pressure from 20-2500 bar (red-to-blue) of RE137 apo (top right), and with 15-bound (bottom left) and 47-bound (bottom right) forms. Selected trajectories are indicated by arrows to show direction of pressure-induced chemical shift changes. All samples contained 195 µM RE137 protein, with 1.2 mM compounds added where indicated.

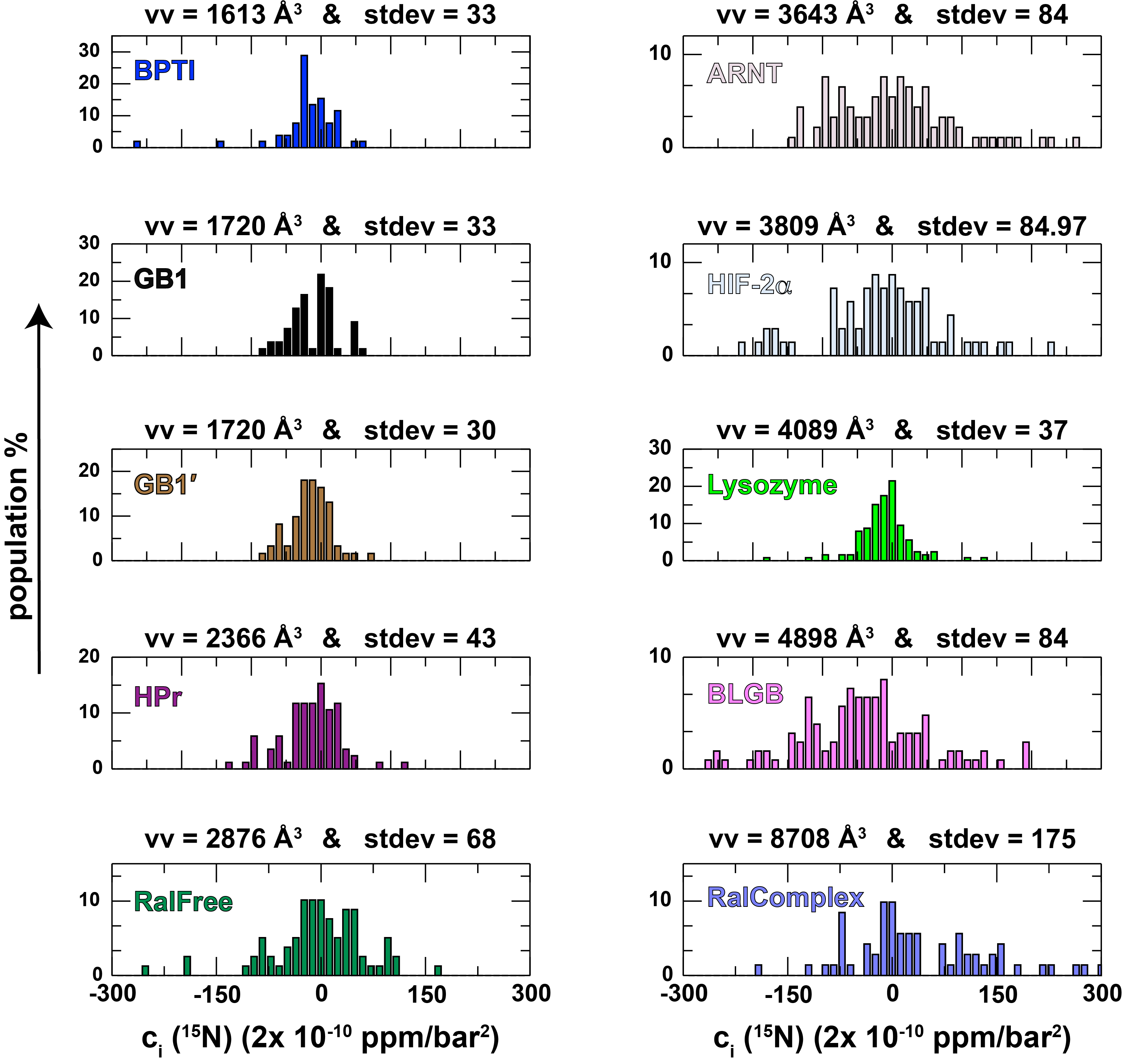

**Fig. S8: Histograms of ^15^N c*_i_* parameters for each of the ten proteins indicated in Figure 3.**

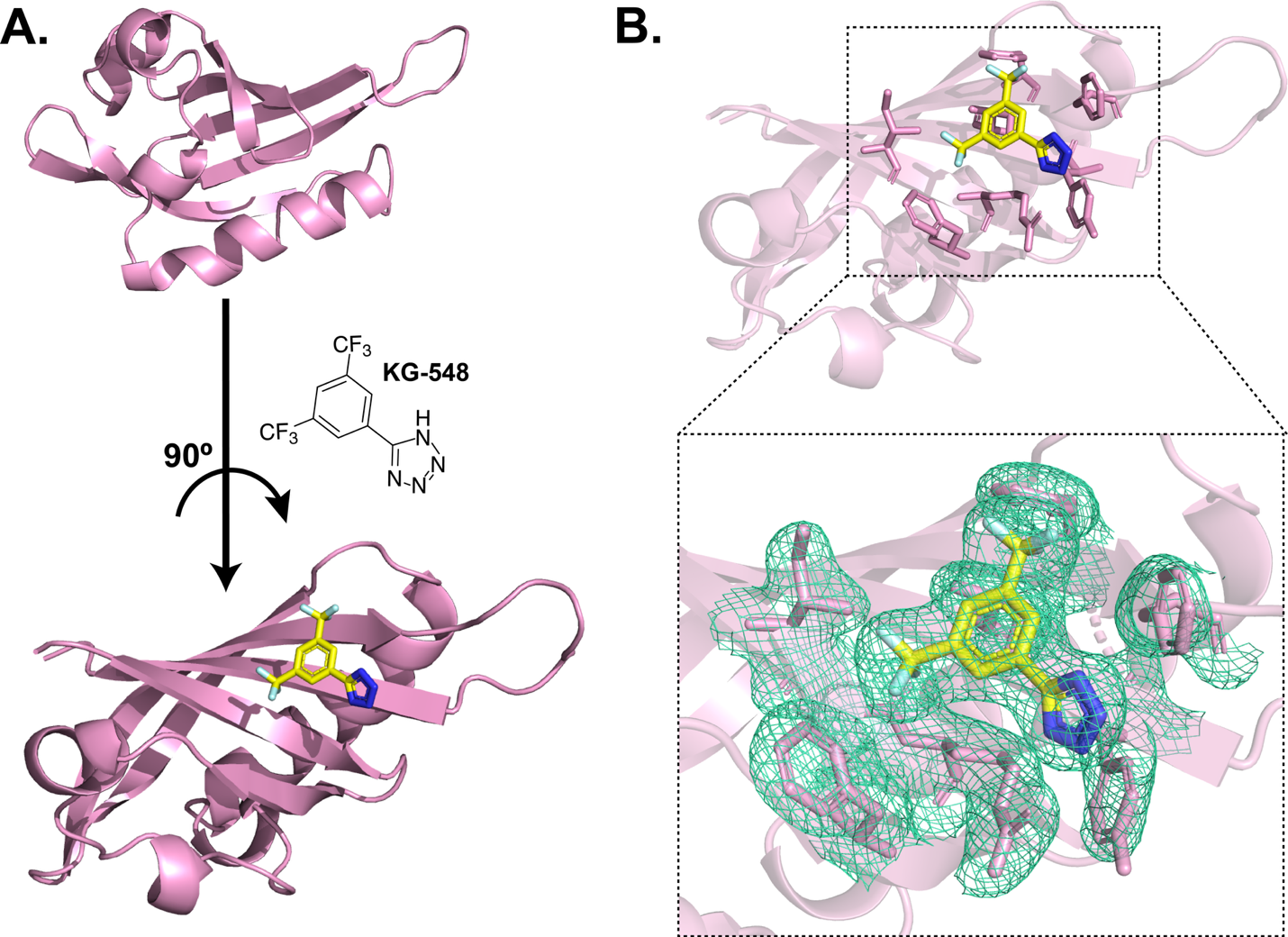

**Fig. S9: X-ray structure of the ARNT PAS-B:KG-548 complex.** (A) X-ray structure of the ARNT PAS-B:KG-548 complex (PDB ID: 84GA) shows KG-548 binding to the exterior beta-sheet surface of the PAS-B domain. (B) Structure showing the stick presentation of residues interacting with KG-548 on the surface of the PAS-B (top) and the electron density map of the residues and KG-548 (bottom).

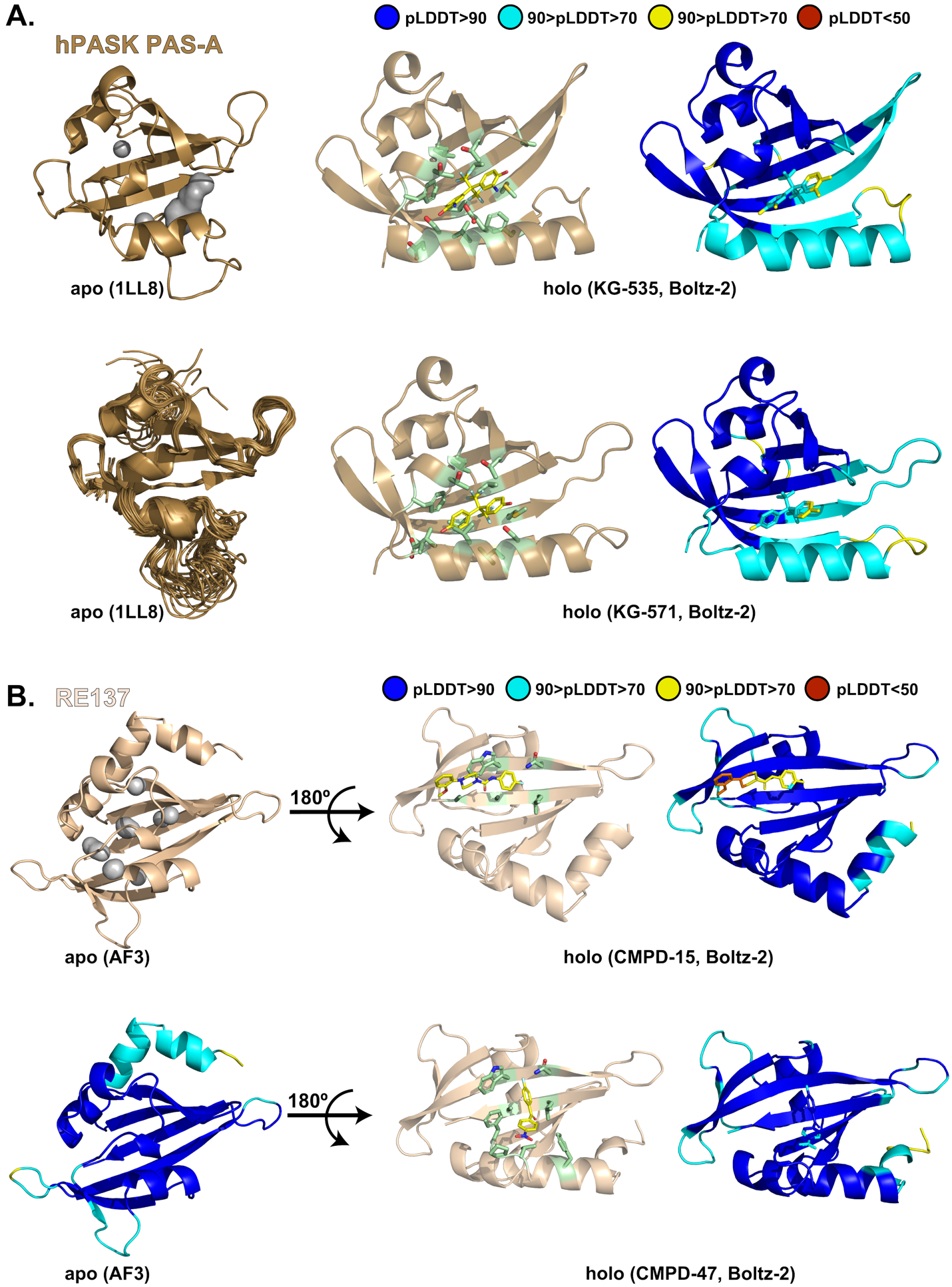

**Fig. S10: Structural models of apo and ligand-bound PAS domains generated using AlphaFold3 and Boltz-2. (A)** hPASK PAS-A: apo structures from the NMR ensemble (PDB ID: 1LL8; representative conformer shown (top left), and multiple conformational states from the NMR ensemble (bottom left) (3). Ligand-bound models with compounds KG-535 (top middle, pLDDT-colored version on the top right) and KG-571 (bottom middle, pLDDT-colored version on the bottom right) were generated using Boltz-2 (23). Predicted pLDDT scores highlight confidence in the ligand-bound poses relative to experimentally observed conformational variability. **(B)** RE137 PAS: apo state predicted with AlphaFold3 (21) (top left, pLDDT-colored version on the bottom left). Ligand-bound models with compound 15 (top middle, pLDDT-colored version on the top right) and compound 47 (bottom middle, pLDDT-colored version on the bottom right) were generated using Boltz-2 (23). Holo models were selected based on pocket-bound poses with minimal steric clash and plausible binding geometry. Binding site residues within 4 Å of the ligand are shown as sticks (ligand carbons in yellow; contacting side chains in pale green). Models are colored by predicted pLDDT scores to illustrate confidence, and holo structures of RE137 are shown with a 180° rotation to visualize cavity access and ligand fit.

**SI References**

1.	T. H. Scheuermann *et al.*, Artificial ligand binding within the HIF2alpha PAS-B domain of the HIF2 transcription factor. *Proc Natl Acad Sci U S A* **106**, 450-455 (2009).

2.	P. B. Card, P. J. A. Erbel, K. H. Gardner, Structural basis of ARNT PAS-B dimerization: Use of a common beta-sheet interface for hetero- and homodimerization. *J Mol Biol* **353**, 664-677 (2005).

3.	C. A. Amezcua, S. M. Harper, J. Rutter, K. H. Gardner, Structure and interactions of PAS kinase N-terminal PAS domain: model for intramolecular kinase regulation. *Structure* **10**, 1349-1361 (2002).

4.	J. R. Huth *et al.*, Design of an expression system for detecting folded protein domains and mapping macromolecular interactions by NMR. *Protein Sci* **6**, 2359-2364 (1997).

5.	F. Corrêa, J. Key, B. Kuhlman, Kevin H. Gardner, Computational Repacking of HIF-2α Cavity Replaces Water-Based Stabilized Core. *Structure* **24**, 1918-1927 (2016).

6.	V. Gonzalez *et al.*, The mosaic structure of the symbiotic plasmid of Rhizobium etli CFN42 and its relation to other symbiotic genome compartments. *Genome Biol* **4**, R36 (2003).

7.	V. Gonzalez *et al.*, The partitioned Rhizobium etli genome: Genetic and metabolic redundancy in seven interacting replicons. *Proceedings of the National Academy of Sciences* **103**, 3834-3839 (2006).

8.	I. Dikiy *et al.*, Diversity of function and higher-order structure within HWE sensor histidine kinases. *J Biol Chem* **299**, 104934 (2023).

9.	S. M. Harper, L. C. Neil, K. H. Gardner, Structural basis of a phototropin light switch. *Science* **301**, 1541-1544 (2003).

10.	P. G. Blommel, B. G. Fox, A combined approach to improving large-scale production of tobacco etch virus protease. *Protein Expr Purif* **55**, 53-68 (2007).

11.	K. L. Cheung *et al.*, Distinct Roles of Brd2 and Brd4 in Potentiating the Transcriptional Program for Th17 Cell Differentiation. *Mol Cell* **65**, 1068-1080 e1065 (2017).

12.	G. Zhang *et al.*, Down-regulation of NF-kappaB transcriptional activity in HIV-associated kidney disease by BRD4 inhibition. *J Biol Chem* **287**, 28840-28851 (2012).

13.	M. Gacias *et al.*, Selective chemical modulation of gene transcription favors oligodendrocyte lineage progression. *Chem Biol* **21**, 841-854 (2014).

14.	M. K. Jang *et al.*, The bromodomain protein Brd4 is a positive regulatory component of P-TEFb and stimulates RNA polymerase II-dependent transcription. *Mol Cell* **19**, 523-534 (2005).

15.	Z. Charlop-Powers, L. Zeng, Q. Zhang, M. M. Zhou, Structural insights into selective histone H3 recognition by the human Polybromo bromodomain 2. *Cell Res* **20**, 529-538 (2010).

16.	Q. Wang *et al.*, Protocols and pitfalls in obtaining fatty acid-binding proteins for biophysical studies of ligand-protein and protein-protein interactions. *Biochem Biophys Rep* **10**, 318-324 (2017).

17.	Y. He *et al.*, Solution-state molecular structure of apo and oleate-liganded liver fatty acid-binding protein. *Biochemistry* **46**, 12543-12556 (2007).

18.	E. Gasteiger *et al.*, ExPASy: The proteomics server for in-depth protein knowledge and analysis. *Nucleic Acids Res* **31**, 3784-3788 (2003).

19.	J. Jumper *et al.*, Highly accurate protein structure prediction with AlphaFold. *Nature* **596**, 583-589 (2021).

20.	R. Evans *et al.*, Protein complex prediction with AlphaFold-Multimer. *bioRxiv* 10.1101/2021.10.04.463034, 2021.2010.2004.463034 (2022).

21.	J. Abramson *et al.*, Accurate structure prediction of biomolecular interactions with AlphaFold 3. *Nature* **630**, 493-500 (2024).

22.	G. Tauriello *et al.*, ModelArchive: A Deposition Database for Computational Macromolecular Structural Models. *J Mol Biol* **437**, 168996 (2025).

23.	S. Passaro *et al.*, Boltz-2: Towards Accurate and Efficient Binding Affinity Prediction. *bioRxiv* 10.1101/2025.06.14.659707, 2025.2006.2014.659707 (2025).

24.	R. J. Quinlan, G. D. Reinhart, Baroresistant buffer mixtures for biochemical analyses. *Anal Biochem* **341**, 69-76 (2005).

25.	B. A. Johnson, From Raw Data to Protein Backbone Chemical Shifts Using NMRFx Processing and NMRViewJ Analysis. *Methods Mol Biol* **1688**, 257-310 (2018).

26.	B. A. Johnson, R. A. Blevins, NMRView: a computer program for the visualization and analysis of NMR data. *J Biomol NMR* **4**, 603-614 (1994).

27.	M. Norris, B. Fetler, J. Marchant, B. A. Johnson, NMRFx Processor: a cross-platform NMR data processing program. *J Biomol NMR* **65**, 205-216 (2016).

28.	E. Koag *et al.*, NMRFx: Integrated Software for NMR Data Processing, Visualization, Analysis and Structure Calculation. *bioRxiv* 10.1101/2025.08.26.672401 (2025).

29.	K. Akasaka, Probing conformational fluctuation of proteins by pressure perturbation. *Chem Rev* **106**, 1814-1835 (2006).

30.	K. Akasaka, H. Li, Low-Lying Excited States of Proteins Revealed from Nonlinear Pressure Shifts in ^1^H and ^15^N NMR. *Biochemistry* **40**, 8665-8671 (2001).

31.	T. H. Scheuermann *et al.*, Allosteric inhibition of hypoxia inducible factor-2 with small molecules. *Nat Chem Biol* **9**, 271-276 (2013).

32.	C. R. Chen, G. I. Makhatadze, ProteinVolume: calculating molecular van der Waals and void volumes in proteins. *BMC Bioinformatics* **16**, 101 (2015).

33.	C. Vonrhein *et al.*, Data processing and analysis with the autoPROC toolbox. *Acta Crystallogr D Biol Crystallogr* **67**, 293-302 (2011).

34.	D. Liebschner *et al.*, Macromolecular structure determination using X-rays, neutrons and electrons: recent developments in Phenix. *Acta Crystallogr D Struct Biol* **75**, 861-877 (2019).

35.	P. Emsley, B. Lohkamp, W. G. Scott, K. Cowtan, Features and development of Coot. *Acta Crystallogr D Biol Crystallogr* **66**, 486-501 (2010).

36.	R. P. Joosten, F. Long, G. N. Murshudov, A. Perrakis, The PDB_REDO server for macromolecular structure model optimization. *IUCrJ* **1**, 213-220 (2014).

37.	P. J. Erbel, P. B. Card, O. Karakuzu, R. K. Bruick, K. H. Gardner, Structural basis for PAS domain heterodimerization in the basic helix--loop--helix-PAS transcription factor hypoxia-inducible factor. *Proc Natl Acad Sci U S A* **100**, 15504-15509 (2003).

38.	T. H. Scheuermann *et al.*, Isoform-Selective and Stereoselective Inhibition of Hypoxia Inducible Factor-2. *J Med Chem* **58**, 5930-5941 (2015).

39.	J. L. Rogers *et al.*, Development of inhibitors of the PAS-B domain of the HIF-2alpha transcription factor. *J Med Chem* **56**, 1739-1747 (2013).

40.	Y. Guo *et al.*, Regulating the ARNT/TACC3 axis: multiple approaches to manipulating protein/protein interactions with small molecules. *ACS Chem Biol* **8**, 626-635 (2013).

41.	M. R. Evans, P. B. Card, K. H. Gardner, ARNT PAS-B has a fragile native state structure with an alternative beta-sheet register nearby in sequence space. *Proc Natl Acad Sci U S A* **106**, 2617-2622 (2009).

42.	T. M. Jacobs *et al.*, Design of structurally distinct proteins using strategies inspired by evolution. *Science* **352**, 687-690 (2016).

43.	A. I. Nash *et al.*, Structural basis of photosensitivity in a bacterial light-oxygen-voltage/helix-turn-helix (LOV-HTH) DNA-binding protein. *Proc Natl Acad Sci U S A* **108**, 9449-9454 (2011).
